# Voluntary Exercise Differentially Affects High-Fat Induced Metabolic Phenotypes Across BXD Mouse Strains

**DOI:** 10.64898/2026.09.28.752290

**Authors:** Arianna Lamanna, Ziyun Zhou, Besma Boussoufa, Emeline Pansart, Pierre Garcia, Tony Heurtaux, Manuel Buttini, Djalil Coowar, Giusi Marcon, Evan G. Williams

**Author notes:** These authors contributed equally (Arianna Lamanna, Ziyun Zhou).

## Abstract

Obesity and metabolic diseases such as diabetes reflect long-term interactions between diet, lifestyle, and genetics – variables that are hard to control in humans but can be systematically varied in mice, though behavioral variables like long-term exercise remain harder to model. Having previously phenotyped ∼50 BXD strains, we identified BXD40, BXD43, and BXD100 as predisposed to voluntary wheel-running (∼6 km/day at 24 weeks) with variable susceptibility to diet-induced obesity and diabetes. We followed these strains from 8–32 weeks on a 60% kcal/fat high-fat (HF) diet, with or without access to running wheels. Wheel usage declined from ∼9 km/day in the first two weeks to a steady ∼5 km/day thereafter yet still improved oral glucose tolerance by 35% over sedentary controls by 24 weeks. Despite equivalent body weight, runners ate 20% more relative to body weight, indicating the glucose benefit was independent of caloric intake.

We next asked whether the gut microbiome contributes to this benefit. Long-term exercise produced only modest, cohort-dependent shifts in cecal microbial composition, though functional profiling revealed exercise-associated differences in carbohydrate- and glucose-metabolism pathways. Reciprocal bedding exchange between sedentary and running BXD43 cages reduced microbial differentiation in a taxon-specific manner and coincided with marked attenuation of the glucose phenotype, despite unchanged running behavior. Together, these findings validate BXD40, BXD43, and BXD100 – alongside the canonical C57BL/6 – as a model for studying diet, exercise, and genetics in metabolic disease, and implicate gut microbial taxa in shaping exercise’s metabolic effects during HF feeding.

## INTRODUCTION

Decades of scientific research have shown a clear, causal connection between excess calorie intake and excess weight gain – often followed by numerous metabolic diseases such as diabetes, hypertension, and nonalcoholic fatty liver disease. The precise contribution and ratios of the different dietary components leading to these effects remains in debate (e.g. the intake of sucrose, saturated fats, carbohydrates, etc.), as is the extent to which moderate exercise mitigates the development of metabolic diseases. In human cohorts, weight loss successfully achieved through dieting is typically reversed within 2-5 years (Beghini et al., 2021; Hall & Kahan, 2018), and while aerobic exercise is indeed beneficial for health, this tends to have only a rather small effect (< 5%) on individuals’ body weights (Oppert et al., 2023). Weight gain – and its strongly-associated metabolic diseases – is not simply driven by a simple, calculable equation between caloric intake versus calories burned, as complex genetic and molecular pathways interact to prevent simple lifestyle changes, such as sustainable diets and exercise regimens, from being sufficient for many people. Due to the long-term nature of metabolic syndrome and the myriad complex and difficult-to-control cofactors, we can use diverse sets of inbred mice strains to determine the effect size of interventions such as diet and long-term exercise on metabolic disease. Over nearly a century of such studies, researchers have identified and created numerous diets which lead to metabolic syndrome in standard inbred laboratory mouse strains, and likewise scientists have identified and created numerous genetic variants which lead to metabolic syndrome, regardless of the intake of a “healthy” or “unhealthy” diet (Delhanty & Visser, 2025). That is, obesity for most individuals comes from a mixture of factors both nature and nurture, referred to as gene-by-environment (GxE) interactions (Herrera-Luis et al., 2024). In natural populations, singular genetic or environmental causes of obesity and metabolic syndrome are rarely observed (Loos & Yeo, 2022), thus it is essential to model the complexity of human metabolism using a controllable model system.

Studies in mice that do not have monogenic causes of obesity (i.e. not *db/db* or *ob/ob* mice) are overwhelmingly performed in the C57BL/6 strain, and they have broadly shown that medium-duration forced exercise on treadmills will induce mice to lose weight and improve glucose tolerance when subjected to *ad libitum* high fat diet (HF) (e.g. 20 minutes/day for a 12 week regiment, (Chang et al., 2020)). However, treadmill experiments also subject mice to significant levels of psychological stress unlike voluntary wheel access (Sasaki et al., 2016), and psychological stress in mice also reduces body weight and improves glucose tolerance (Hatton-Jones et al., 2022). To avoid this confounding factor, studies on exercising mice have provided voluntary access to running wheels over similar 12-week periods, yet these studies have had conflicting results: some show that voluntary exercise *does* mitigate weight gain on HF (Hicks et al., 2016) while others show that it does not (Jung & Luthin, 2010). As we know that genetic background and the amount of exercise play a role in the development of obesity and diabetes, we hypothesized that other inbred strains may provide new insight into these interactions and their effects. We have previously performed metabolic phenotyping on around 50 different recombinant inbred strains from the BXD murine genetic reference population (Williams et al., 2016). These strains have wide variance in their natural propensity to exercise, their body weight, susceptibility to HF-induced obesity and glucose intolerance, and many other key parameters such as expected lifespan (Roy et al., 2021). While mice, which voluntarily exercised, could reverse some of the weight gain induced by HF, our earlier study only permitted 10 days of running activity, before mice were returned to standard (wheel-less) housing (Williams et al., 2016). Mice are widely known to have rapidly diminishing usage of running wheels after around 2 weeks, often dropping to a steady rate of around 1/3^rd^ their initial use when the wheel is first introduced (Jung & Luthin, 2010).

One additional confounding factor in using mice to observe the contribution of voluntary exercise to metabolic effects is that an exercise wheel may not be strictly necessary for exercise: mice also have sufficient space in a normal housing cage to get low-intensity exercise by simply walking around. Developments in home cage monitoring (HCM) over the past 15 years have led to the implementation of mouse housing that can record mice without the necessity of surgically implanted tracking devices. Early such technologies used invisible laser beam grids that recorded animal movement, but with limitations in that normal housing material, such as bedding/nesting, could disrupt the sensors (Voikar & Gaburro, 2020). More recent HCM devices, such as the Tecniplast DVC used in this study, allow animal activity to be tracked via perturbations in electromagnetic sensors caused by animals’ movements (Pernold et al., 2019). These newer systems allow animals to be tracked without implants and in completely standard housing, including normal bedding and enrichments. Thus, it is now possible to continuously record the typical activity of mice throughout their lives.

In this study, we have followed 116 individual mice from 3 BXD strains: BXD40, 43, and 100, on *ad libitum* 60% kcal/fat (5.1 kcal/g) HF from 8 to 32 weeks of age, with cohorts in cages with access to monitored running wheels, or in HCM without access to wheels. Food intake and weight were monitored weekly, and oral glucose response was measured at 24 weeks of age. While both genetic background and exercise played a role in glucose response, we observed that in these lines, increased physical exercise led to an increased caloric intake, and little impact on body weight —although exercise did substantially improve glucose response. Identifying and studying these differences is key to understanding how exercise affects obesity and related fitness parameters in humans: these strains are different from one another and from C57BL/6, so are humans different from one-another. These three strains, which we have previously phenotyped in short-duration exercise, demonstrate models that are suited for identifying the diversity of cellular mechanisms and responses driving the variety of outcomes in genotype, diet, and exercise-dependent traits that occur across populations (Andreux et al., 2012).

Given the growing evidence that the gut microbiome contributes to host energy metabolism and glucose homeostasis (Cani et al., 2019) , we additionally investigated whether exercise-associated alterations of the gut microbiome could contribute to the metabolic effects observed in running mice. Physical activity has previously been associated with changes in gut microbial composition and functional potential, including under conditions of diet-induced obesity, although the magnitude and direction of these effects vary considerably between experimental settings (Allen et al., 2015; Evans et al., 2014; Zhang et al., 2026). Importantly, microbial composition is also strongly influenced by diet and by host genetic background (Perez-Munoz et al., 2019; Zhou et al., 2026). We first used cecal shotgun metagenomic sequencing to characterize exercise-associated differences in microbial taxonomic composition and functional potential across independent experimental cohorts. Exercise-associated taxonomic differences were relatively modest and varied across cohorts, with *Lachnospiraceae* identified as the only significantly different bacterial family in Cohort 1, an effect that was not reproduced in subsequent BXD43 cohorts, but which is consistent with a previous study con diet (Zhou et al., 2026). Despite this limited and cohort-dependent taxonomic response, functional profiling identified exercise-associated differences in several microbial pathways, particularly those involved in carbohydrate and energy metabolism.

To further investigate the potential contribution of the gut microbiome to exercise-associated metabolic effects, we established an additional BXD43 bedding-exchange cohort (Cohort 5) in which soiled bedding was reciprocally transferred between sedentary and running cages on a weekly basis throughout the whole duration of the study. In this cohort, fecal samples were collected longitudinally and analyzed by shotgun metagenomic sequencing at seven selected time points (weeks 8, 9, 11, 13, 16, 19, and 31), allowing us to follow microbiome dynamics over the course of HF feeding, voluntary exercise, and reciprocal bedding exchange. This design enabled us to assess whether sustained microbial exchange reduced microbiome differences between sedentary and running mice and whether this was accompanied by an attenuation of their metabolic differences rather than using a conventional fecal microbiota transplantation approach based for example on oral gavage, we selected repeated reciprocal bedding exchange to provide continuous and non-targeted microbial exposure throughout the intervention. Mice are naturally coprophagic, and transfer of used bedding containing fecal material can therefore facilitate the horizontal transmission of gut microorganisms between animals. Soiled-bedding transfer has also been used following an initial fecal microbiota transplantation by oral gavage to provide continued exposure to donor-derived fecal material and thereby sustain microbial transfer over time (Gheorghe et al., 2021; Stebegg et al., 2019). In addition, dirty bedding transfer has been applied experimentally to reduce cage-associated microbiome variation and homogenize microbial communities between groups (Miyoshi et al., 2018). In contrast to oral gavage, repeated reciprocal bedding exchange was intended to generate a continuous and relatively uncontrolled microbial exchange between sedentary and running animals, more closely resembling the passive horizontal exchange of microorganisms that occurs between individuals sharing an environment. This was particularly relevant given evidence that gut microbial communities can contribute to obesity-associated metabolic phenotypes and that transplantation of obesity-associated microbiota can transmit aspects of these phenotypes to recipient mice (Leocadio et al., 2019). Moreover, co-housing experiments have demonstrated that microbial exchange between mice can modify microbiota-associated metabolic phenotypes (Ridaura et al., 2013), supporting our hypothesis that microbial transfer through bedding exchange between Sed and Run mice could reduce exercise-associated microbiome differences and potentially attenuate the associated differences in metabolic phenotype.

As a complementary pilot experiment, we performed reciprocal bedding exchange between sedentary BXD43 mice maintained on either control diet (CD; *n* = 6) or HF diet (*n* = 7). Because diet is a major determinant of gut microbial composition, this comparison was included as a positive control to assess whether bedding exchange could promote microbial exchange between groups with initially more distinct microbiomes. Despite the strong diet-associated differences, microbial profiles showed evidence of convergence following bedding exchange, supporting the effectiveness of this approach in promoting microbial transfer. We then examined the BXD43 sedentary and running mice, in which longitudinal fecal metagenomics revealed dynamic, time-dependent changes in microbial composition throughout the bedding-exchange intervention, while endpoint cecal differences between exercise conditions were relatively limited. We next asked whether this reduced microbial differentiation was accompanied by changes in the metabolic phenotype. Notably, the pronounced difference in glucose response between sedentary and running BXD43 mice observed in Cohorts 1–4 was strongly reduced in Cohort 5.

Together, these observations show that reciprocal bedding exchange can promote microbial convergence, between groups fed with distinct diets and was accompanied by reduced microbiome and glucose-response differences between sedentary and running mice. While these findings are consistent with a potential contribution of microbial exchange to the metabolic phenotype, they remain associative and exploratory and do not establish a causal microbiome-mediated effect.

## RESULTS

### Highly Heritable Fitness Parameters Used to Select BXDs 40, 43, and 100 For Long-Term Exercise Phenotyping

To select the strains used for this study, we re-examined the data from our large BXD screening study (Williams et al., 2016), where 57 strains were phenotyped on either Harlan Chow (CD) or Harlan 06414 60% HF diets from 8 to 29 weeks of age, with 10 days of access to a running wheel from 22-24 weeks of age while solitarily housed. We have observed that the heritability (h^2^) of these key metabolic traits is very high: body weight h^2^=0.76, voluntary exercise h^2^=0.64, and oral glucose tolerance test response h^2^=0.82. Given the high heritability of these traits, we expected to be able to reproduce and build on these prior findings. To do so, we selected three strains – BXD40, 43, and 100 – which had radically different propensity to develop obesity and metabolic disease when fed with HF (**Figure 1A and Figure S1D**). BXD40’s body weight is nearly unaffected by HF (27 vs 32g), while BXD43 is lean on CD while obese on HF (33g vs. 50g), and BXD100 is large on CD and obese on HF (42g vs. 54g). Furthermore, all three strains were excellent runners: BXD100 runs around 4 km/night, BXD40 6.5 km/night, and BXD43 7.7 km/night (**Figure 1B**), and diet had no overall effect on exercise wheel usage. The mice in this study only had 10 days of access to a running wheel over 22 weeks of HF, thus HF-fed BXD40s did not remain thin due to their wheel usage, so we first hypothesized that differences in normal home cage movement behavior (monitored over 48 hours at 24 weeks of age) may be compensating. However, both BXD40 and BXD43 were exceptionally active, while BXD100 (as with its wheel usage) was only somewhat more active than the average strain (**Figure 1C**). Despite these pronounced exercise traits, all three strains had very average glucose responses in both dietary conditions, with a mild propensity for HF-induced glucose intolerance at 17 weeks of age, after 11 weeks of HF feeding (**Figure S1A**). We also previously observed that 10 days of running can reduce body weight by around 10% in both dietary cohorts (**Figure S1B-C**). With these prior results in mind, we designed a study to allow us to continuously monitor normal movement in a “sedentary” HCM cohort, while an active “running” cohort would have *ad libitum* access to a recorded running wheel, on the same 60% kcal/fat HF, and over a similar range of ages (8-32 weeks old) (**Figure 1D**). In this study, we measured body weight and food intake weekly during this period, and the results of an intraperitoneal glucose tolerance test (IPGTT) at 24 weeks of age. We hypothesized that mice resistant to the negative effects of HF may not be directly genetically resistant to the diet but instead may continue to exercise by running around their cage –albeit likely less than for mice with access to a running wheel.

**Figure 1.**
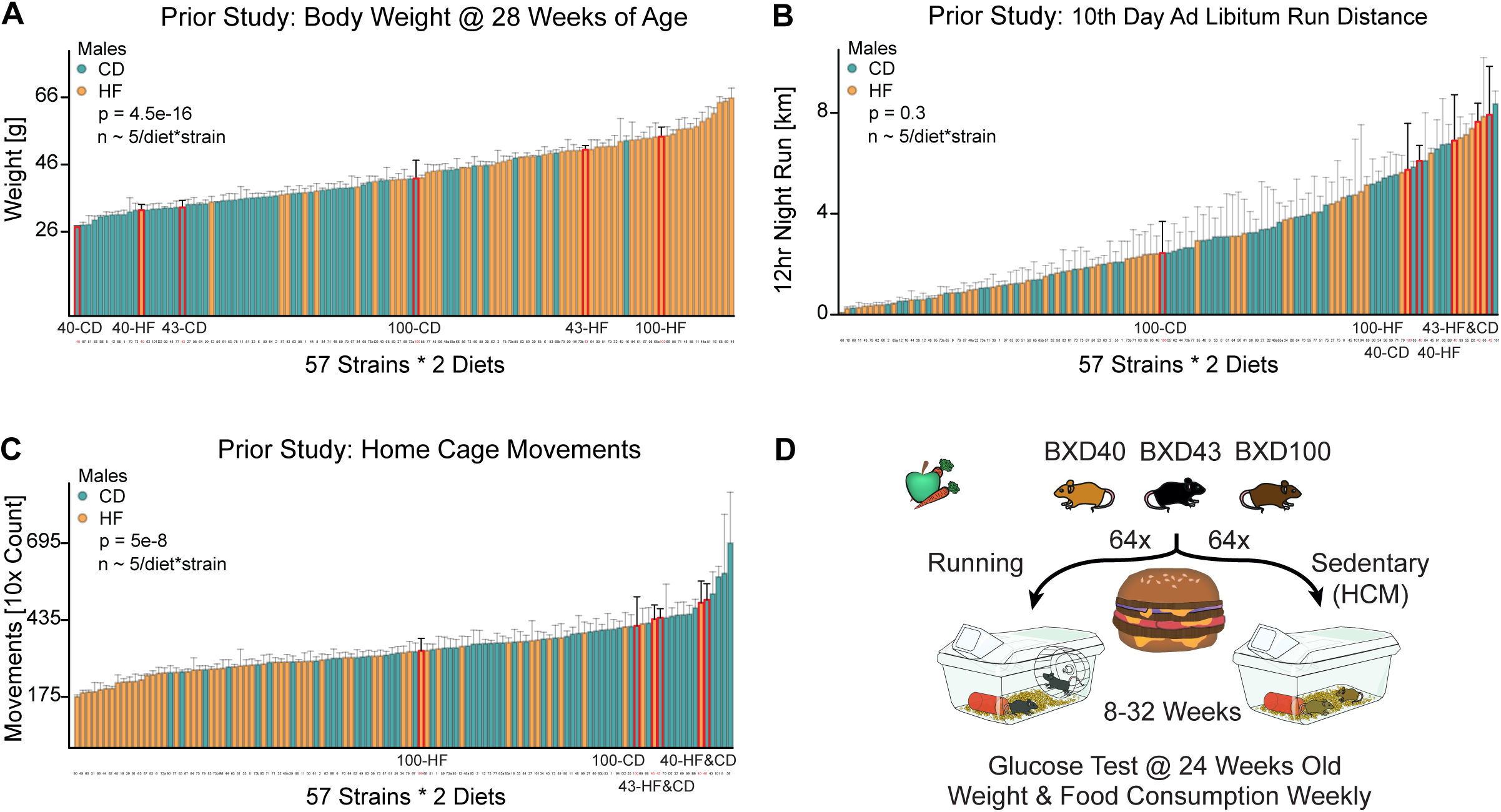
BXD Strain Selection, Hypothesis Development, and Study Design (**A**) Body weight at 28 weeks of age (20 weeks of CD or HF feeding) in a prior study on 57 BXD strains. (**B**) The final cumulative daily run for a solitary mouse with an exercise wheel after nine days of habituation, in the same BXD cohort at around 24 weeks of age. (**C**) Home cage monitoring over 24 hours for a solitary mouse after one day of habituation, in the same cohort. (**D**) Study design using a subset of the previous strains on HF with continuous access to a running wheel or HCM activity monitoring (Cohort 1-4).

**Figure S1.**
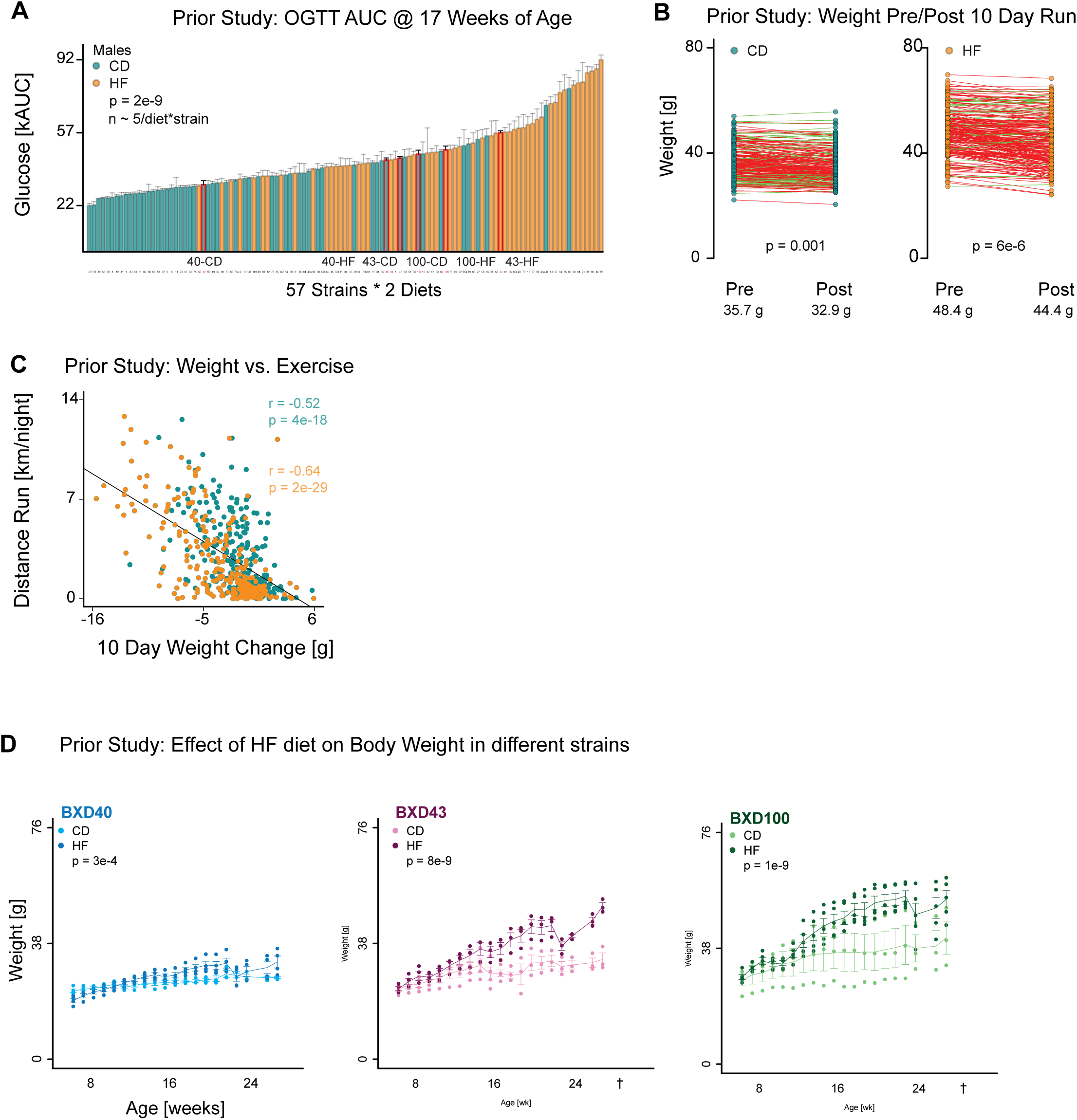
Previous Results and Knowledge About BXD Metabolic Phenotypes (**A**) Oral glucose response in a prior study on 57 BXD strains. (**B**) Difference in body weight before and after 10 days of voluntary exercise in a prior study on 57 BXD strains. (**C**) Bodyweight and Distance Run association in CD and HF feed BXD mice. (**D**) Body weight to HF response over time in BXD40, BXD43 and BXD100 mice.

### Voluntary Exercise Only Slightly Improves Glucose Response and Lowers Body Weight Gain in HF Cohorts

While we knew from our prior 10-day exercise study that the BXD40, 43, and 100 (as well as C57BL/6) will use a running wheel voluntarily, we also knew from prior literature that C57BL/6 mice reduce their activity wheel usage over time, typically dropping by half from around ∼10 km/day at ∼10 weeks of age to ∼5 km/day by 20 weeks of age (Figueiredo et al., 2009). Thus, we first needed to validate that our strains would continue to voluntarily exercise throughout the duration of the experiment. We indeed observe that average weekly running distance dropped by 60% from 64 to 25 km/week/cage between 9 weeks versus 29 weeks of age (p=3e-6; **Figure 2A**). This decline in running is consistent for nearly all running mice: of the 46 running mice, all but 2 reduced their median running activity in the first 10 weeks of the study, compared to their median running in the following 10 weeks, dropping by 43% (p = 2e-4; **Figure 2B**). Due to space and equipment limitations, only 16 mice could be run per cohort, but we observed no difference in running distance between the three cohorts (ANOVA, p = 0.16, **Figure 2C**). Compared to the decrease in activity wheel usage over time, we expected a less significant effect in normal home cage over time, based on prior literature showing only a slight decline with time (Pernold et al., 2021). Indeed, we observed that our sedentary cohorts moved only 27% less between 14 weeks to 29 weeks of age (p = 0.01, **Figure 2D**). (Note that 14 weeks is used as the baseline for HCM, as the first 6 weeks of DVC data were lost for Cohort 1.) This decline in movement at the young age also quickly stabilized, with mice not having any further decline in HCM activity between 17±3 weeks to 26±3 weeks of age (p = 0.51, **Figure 2E**). In contrast to the activity wheel, we did observe differences across cohorts for their overall home cage activity, with a significant elevation in Cohort 3, which moved nearly 50% more in HCM than did the first two cohorts (p < 2e-16, **Figure 2F**).

**Figure 2.**
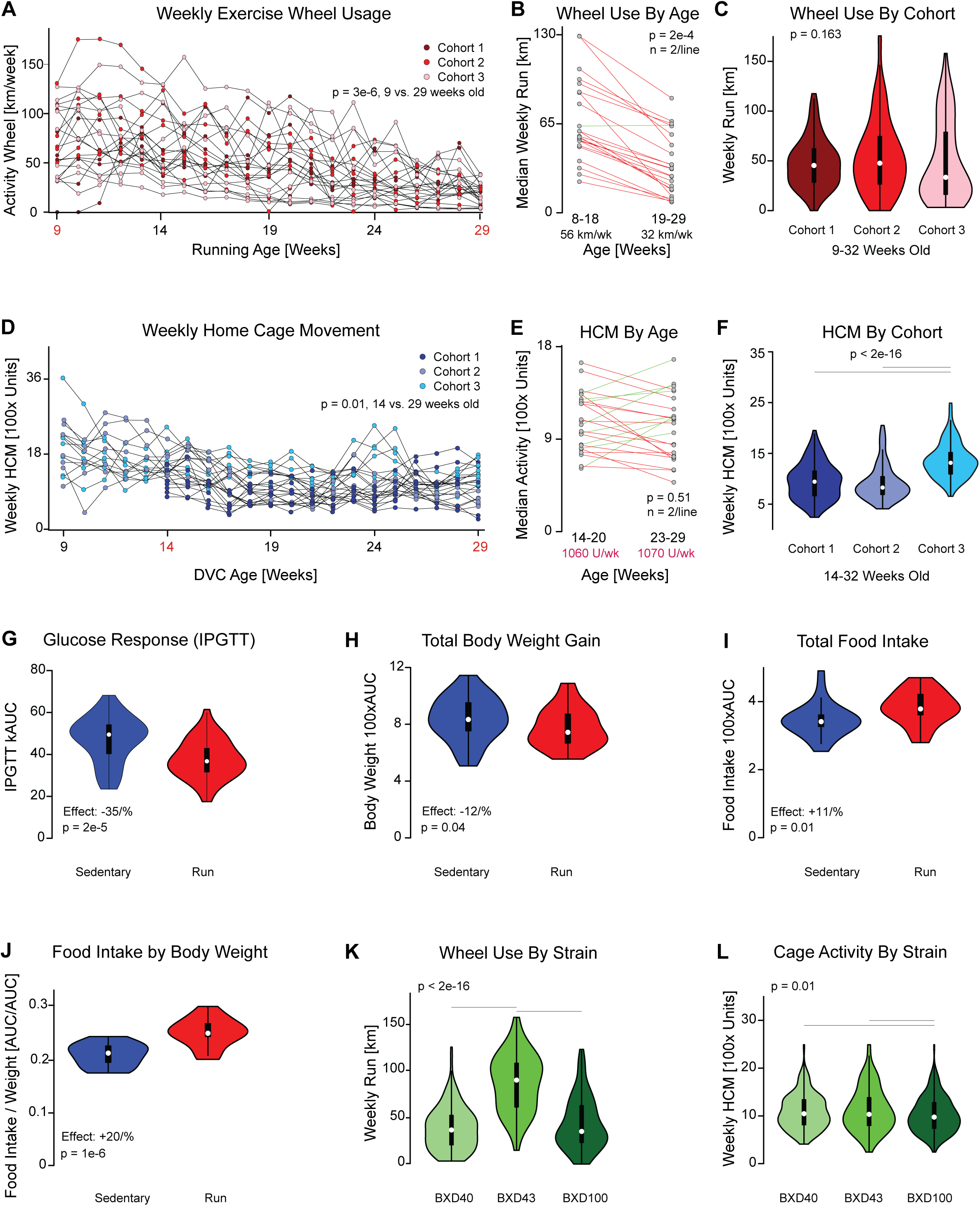
Exercise and Home Cage Monitoring in the BXDs (**A**) Weekly total activity wheel usage for all running cohorts across time. (**B**) Slopegraph comparison of median movement between weeks 8-18 versus 19-29 for all running cohorts. (**C**) Violin plot of running distance for all mice in each cohort. (**D**) HCM total activity for all sedentary cohorts across time. (**E**) Slopegraph comparison of median movement between weeks 8-18 versus 19-29 for all sedentary cohorts. (**F**) Violin plot of HCM activity for all mice in each cohort. (**G**) IPGTT glucose response decrease of 35% in running cohorts vs. sedentary cohorts. (**H**) Body weight decreased by 12% in running cohorts vs. sedentary cohorts. (**I**) Running mice ate around 11% more than sedentary mice, (**J**) or around 20% more than sedentary mice if corrected for the slightly decreased body weight of running mice. (**K**) BXD43 ran more than twice as much as the other two strains, but (**L**) HCM activity was roughly constant across all three lines (6% decrease for BXD100).

**Figure S2.**
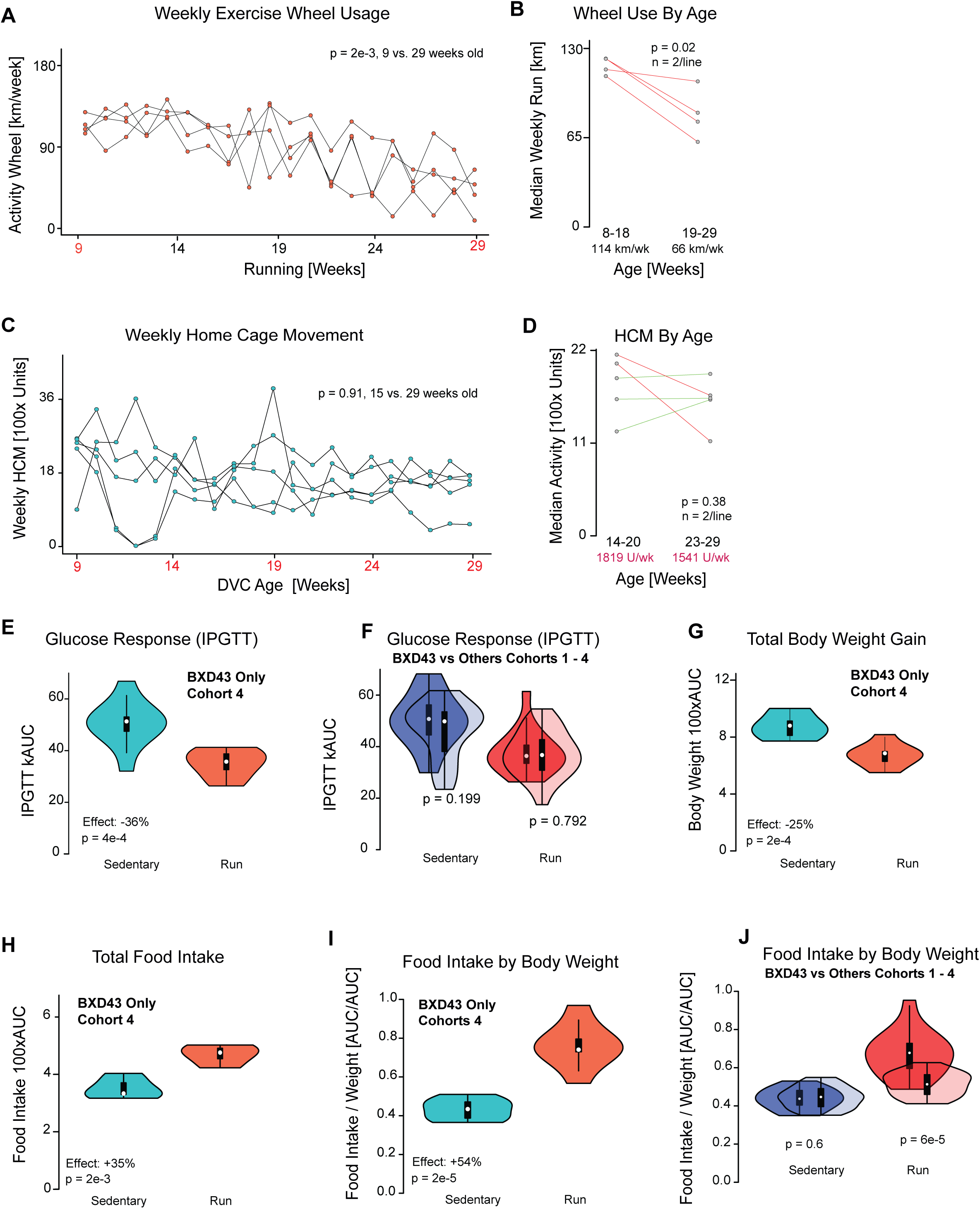
BXD43: Cohort 4: Validation (**A**) Weekly total activity wheel usage for cohort 4 females over time. (**B**) Slopegraph comparison of median movement between weeks 8-18 versus 19-29 for all running cohorts. (**C**) HCM total activity for all sedentary cohorts across time. (**D**) Slopegraph comparison of median movement between weeks 14-20 versus 23-29 for all sedentary cohorts. (**E**) IPGTT glucose response decrease of 38% in running cohorts vs. sedentary cohorts. (**F**) Body weight decreased by 22% in running cohorts vs. sedentary cohorts. (**G**) Running mice ate around 31% more than sedentary mice, (**H**) or around 68% more than sedentary mice if corrected for the slightly decreased body weight of running mice.

Given that the mice exercised, we expected to identify significant changes in the weight gain and early indicators of metabolic syndrome, such as hyperglycemia during the IPGTT. Indeed, during the IPGTT experiment at 24 weeks of age, sedentary mice had at least a 35% worse glucose response curve than the running cohort (p = 2e-5, **Figure 2G**). (note, this difference is likely higher, but the glucometer used has a maximum value of 600 mg/dL, which was reached at 30 minutes by 8 running mice, compared to 24 sedentary mice (**Table S1**)). We next checked the overall change in body weight from 8 to 32 weeks of age (**Table S3**) and observed only a 12% increase in body weight for sedentary mice compared to running cohorts (p = 0.04, **Figure 2H**). This was initially a surprise, given the significant use of the running wheel, and we hypothesized that the running mice may have increased their food intake. This was indeed the case, and running mice ate about 11% more than the sedentary cohort (p = 0.01, **Figure 2I, Table S4**). Factoring in the smaller size of the running mice, we observed that the running mice ate 20% calories compared to their body weight versus the sedentary mice (p = 1e-6, **Figure 2J**), in line with expectations from literature (Kelly et al., 2011).

While *ad libitum* voluntary exercise led to a significant, although not massive, improvement in metabolic fitness, we expected that genetic background would play a role in these outcomes. For instance, BXD43 ran a median of 90 km/week throughout the experiment, far exceeding the average of the other strains combined median of 38 km/week (p = 1e-16, **Figure 2K**). Conversely, home cage activity of all three strains was nearly equivalent, with only BXD100 being very marginally less active than the other two strains (−6%; p = 0.01, **Figure 2L**). Thus, we overall confirmed that BXD43 is a highly active mouse model for voluntary exercise, and that the high exercise wheel usage does not correspond to a higher basal activity in the absence of a wheel. We thus selected three avenues of analysis to further pursue this finding. First, do most active running mice perform metabolically better than the least active, and is there any connection between HCM activity and metabolic fitness? Second, by testing a fourth cohort using only BXD43, to increase the sample size and replicate the main finding, do we confirm that strain’s particular connection between exercise, glucose response, weight, and food intake? Finally, can we identify cofactors which indicate the causal mechanisms, strength, or predictive variables connecting genetic background, diet, and exercise to metabolic disease outcomes?

### Increased Activity Does Not Strongly Predict Improved Metabolic Outcomes

We have identified how genetic predisposition (**Figure 1A–C**) and voluntary exercise (**Figure 2G–J**) play a significant role in modulating the metabolic effects of HF diets. Human cohort studies have demonstrated an approximately linear inverse dose–response relationship between physical activity and the incidence of type 2 diabetes, with progressively higher activity levels associated with a lower risk of disease development (Strain et al., 2023).

In this study, we observe a ∼4-fold range between the least active (∼400 km run) and most active (∼1600 km run) runners (**Figure 3A**), and a similar ∼3-fold range between the least and most active sedentary cohorts with HCM (**Figure 3H**). We observed that total weekly running distance was positively associated with the food intake to body weight ratio (r = 0.60, p = 4e-3; **Figure 3C**). In contrast, neither food intake nor body weight alone showed a significant association with running distance (**Figure 3 A–B)**, likely because body weight is strongly influenced by food intake independently of exercise. No significant associations between Total DVC activity, body weight and food intake have been shown in the sedentary cohort (**Figure 3H-J**).

**Figure 3.**
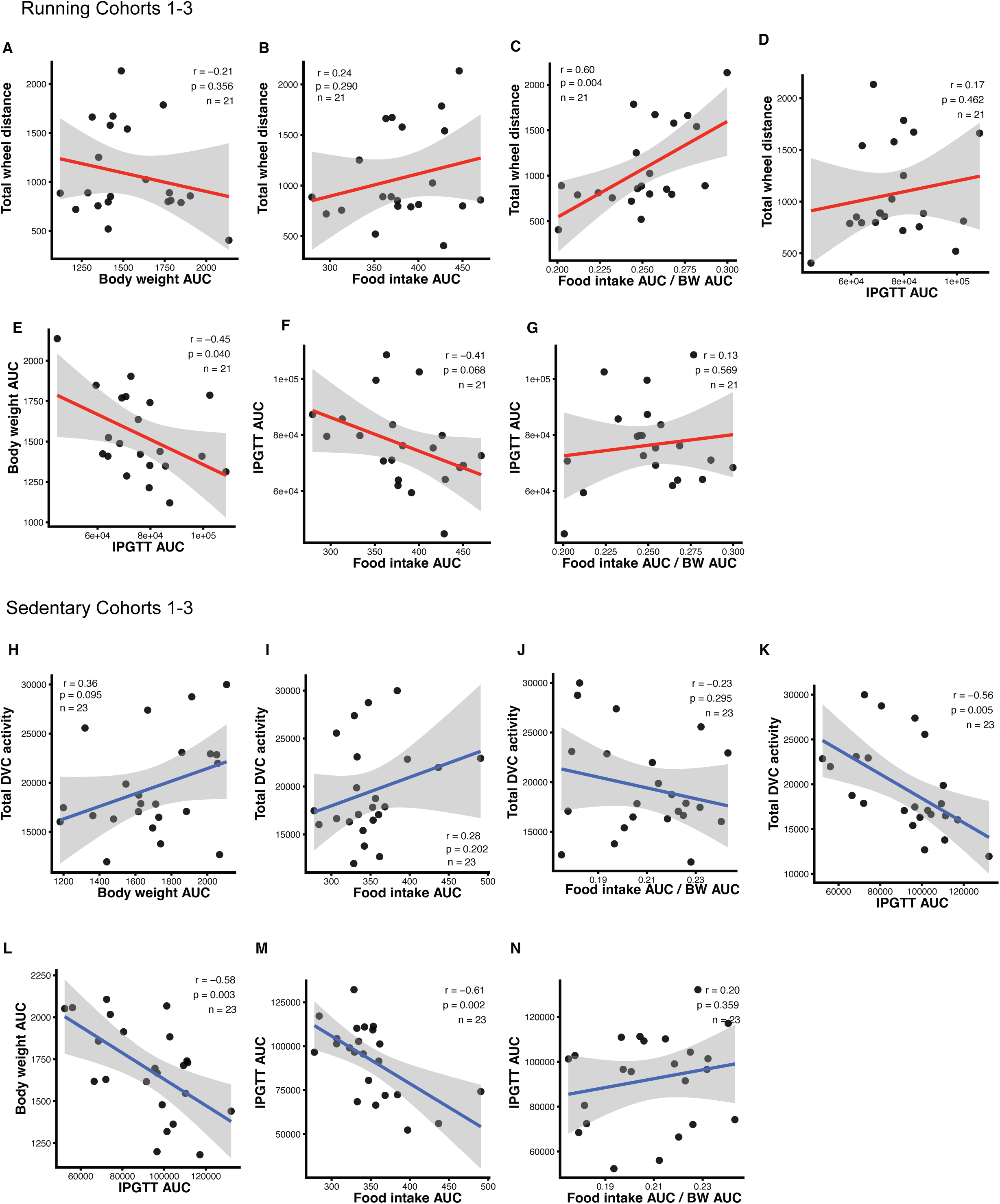
Activity and Metabolism: Associations Within Sedentary and Running Cohorts Associations between activity and metabolic parameters were assessed separately in running and sedentary mice from Cohorts 1–3. For running mice, total wheel-running distance was correlated with (**A**) body-weight AUC, (**B**) food-intake AUC, (**C**) food intake-to-body-weight AUC ratio, and (**D**) IPGTT AUC. (**E**) Body-weight AUC was negatively associated with IPGTT AUC, while IPGTT AUC was further compared with (**F**) food-intake AUC and (**G**) food intake-to-body-weight AUC ratio. For sedentary mice, total DVC activity was correlated with (**H**) body-weight AUC, (**I**) food-intake AUC, (**J**) food intake-to-body-weight AUC ratio, and (**K**) IPGTT AUC. (**L**) Body-weight AUC was negatively associated with IPGTT AUC, while IPGTT AUC was also negatively associated with (**M**) food-intake AUC and (**N**) food intake-to-body-weight AUC ratio. Pearson’s correlation coefficients (r) and p-values are shown in each panel.

**Figure S3.**
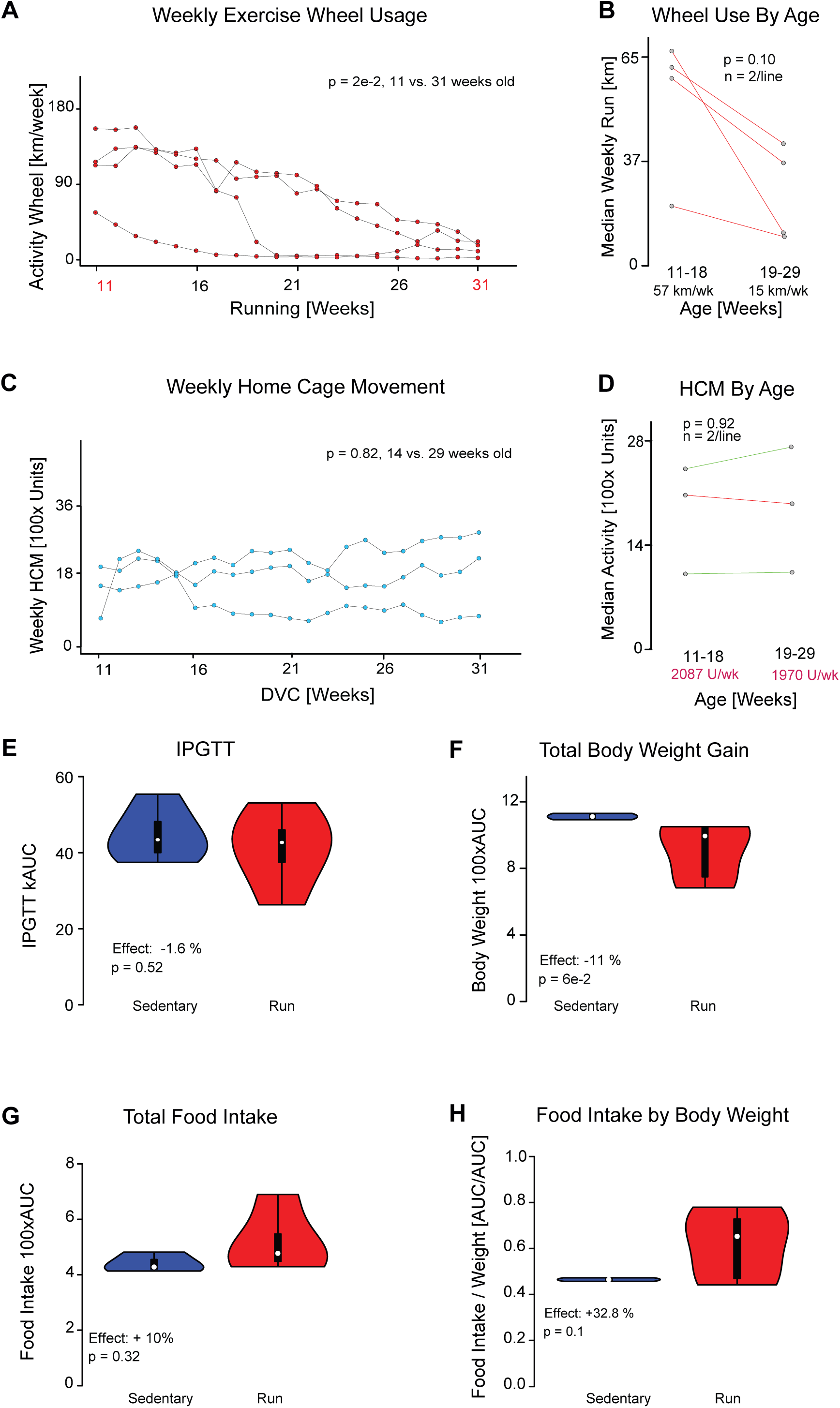
Cohort 4 Male Mice only (**A**) Weekly running-wheel activity across the experimental period, showing a significant decline with age (p = 2e-2). (**B**) Comparison of median weekly running distance between weeks 11–18 and 19–29 (p = 0.10). (**C**) Weekly home-cage monitoring (HCM) activity across the experimental period, which remained stable over time (p = 0.82). (**D**) Comparison of median HCM activity between weeks 11–18 and 19–29 (p = 0.92). (**E**) IPGTT glucose response was 1.6% lower in Run compared with Sed mice (p = 0.52). (**F**) Body-weight gain was 11% lower in Run mice (p = 6e-2). (**G**) Total food intake was 10% higher in Run mice (p = 0.32), and (**H**) the food intake-to-body-weight ratio was 32.8% higher in Run mice (p = 0.10).

Interestingly, the glucose response during the IPGTT was inversely associated with body weight in both sedentary (r = -0.58. p = 3e-3, **Figure 3L**) and running mice (r = -0.45. p = 4e-2, **Figure 3E**), indicating that mice with higher body weight unexpectedly exhibited a lower glucose response. Although increased body weight and adiposity are generally associated with impaired glucose homeostasis, these parameters are not invariably coupled. Indeed, (Yao et al., 2024) reported that HFD-fed mice with greater body weight and adiposity paradoxically exhibited improved glucose tolerance, demonstrating that increased body weight does not necessarily translate into impaired glucose handling. Similarly, Douglass et al. (2023) demonstrated that glucose tolerance can be regulated independently of body weight through hypothalamic microglial signaling, highlighting the existence of mechanisms capable of uncoupling glucose homeostasis from adiposity.

Such mechanisms may be particularly relevant in the context of physical activity. Voluntary wheel running has recently been shown to improve glucose tolerance and insulin sensitivity in HFD-fed mice while simultaneously suppressing hypothalamic inflammation and microglial activation, with some of these effects occurring independently of changes in body weight (Sasaki et al., 2024). Exercise can additionally enhance skeletal muscle glucose uptake (Sylow et al., 2017), providing another mechanism through which physical activity may improve glucose metabolism independently of body weight. Therefore, exercise-induced adaptations may contribute to the unexpected inverse association between body weight and glucose response observed in our mice. Consistent with this hypothesis, the glucose response was also inversely associated with total DVC activity in sedentary mice (r = -0.56, p = 5e-3, **Figure 3K**), indicating that animals displaying greater spontaneous physical activity exhibited better glucose tolerance. However, this was not the case for the running cohort (**Figure 3D**).

In sedentary mice, IPGTT response was negatively associated with food intake (**Figure 3M**), whereas no such association was observed in the running cohort (**Figure 3F**). However, when food intake was normalized to body weight, no significant association with IPGTT response was observed in either group (**Figure 3G**, **Figure 3N**).

Together, these findings suggest that the relationship between body weight and glucose homeostasis is complex and may be influenced by physical activity, genetic background, body composition, and weight-independent mechanisms of glucose regulation.

### Validation Cohort: Expansion & Re-Testing of BXD43

Given that BXD43 had the most striking activity wheel usage in both our original large-scale screening study and in this long-duration exercise cohort (i.e. **Figure 1A**, **Figure 2K**), we decided to test this strain again, using all of the available running wheels (i.e. testing 32 BXD43 mice at once). We observed again that this strain was highly active, with the four cages of BXD43s running 114 km/week on the first full week of the experiment (i.e. 9 weeks of age, **Figure S2A**). As with the previous cohorts, exercise use dropped by about half during the first and second halves of the study period, down to around 66 km/week (p= 2e-3, **Figure S2B**). Notably, while exercise dropped equivalently, the BXD43 remained more active when comparing individual times to the other strains: pairs of BXD43s at 8 months of age ran the same ∼60 km/day/mouse as did the other cohorts at 3 months of age (i.e. **Figure 2B**). As with the first three cohorts, HCM did not significantly change over time (**Figure S2C**), with equivalent movement in both halves of the experimental time (**Figure S2D**). However, it is worth noting that just as BXD43 ran roughly twice as much with the exercise wheels, the strain was also ∼1.6x more active than the other strains with respect to home cage movement (i.e. comparing **Figure S2D** with **Figure 2E**). As with the previous cohorts, exercising mice had substantially better glucose tolerance responses (36% improved, p = 4e-4, **Figure S2E**). However, there was no difference between the BXD43s and the BXD40s and BXD100s from the previous Cohorts (**Figure S2F**). The Cohort 4 running mice had more significantly diminished body weight (−25%, p = 2e-4, **Figure S2G**) compared to the first three cohorts, despite an even larger increase in food intake (+35% p = 2e-3, **Figure S2H** in BXD43, versus a +11% increase in the other cohorts), particularly when adjusted to body weight (+54%, p = 2e-5, **Figure S2I**, versus a +20% increase in the other cohorts). In fact, the food intake by body weight ratio is significantly higher in the running BXD43 compared to the other strains (Cohorts 1-3) of the homonymous group (p= 6e-5**, Figure S2J**).

As the initial cohorts were all tested exclusively in female mice, we also aimed to test BXD43 in males. Four pairs of males were placed in cages identically as to the female cohorts and followed for the same period. While running activity did drop over time from (p = 2e-2, **Figure S3A**) the variance in cage-to-cage and week-to-week was highly elevated (**Figure S3B**). As with all other cohorts, HCM activity was essentially constant over time (**Figure S3C-D**). Unlike the female cohorts, we observed no difference in glucose response nor body weight in the exercising versus sedentary mice (**Figure S3E-H**). However, it is important to note a major caveat for the males: three of the four males in running cages severely injured their littermate to the extent that it required euthanasia, as did two of the four males in the sedentary cages. Even prior to these severe injuries, the male cages had far more dramatic dominant-versus-submissive animals in the running cages, with ∼15 g differences in weight between the males for all three running cages that had the severe fighting. Consequently, the variation of each cage was high and running wheel usage may not have been even approximately distributed between the two males per cage. While these challenges with male aggression may be addressable with a redesign of the study, the current male cohort was not considered suitable for future analysis, and we chose to focus on the female cohorts for molecular analysis.

### Causal Associations and Possible Molecular Mechanisms and Variables Predicting Metabolic Health Outcomes

We have seen that these three highly active strains are able to mitigate, although by no means eliminate, the negative metabolic effects of *ad libitum* HF feeding. We next sought to determine whether voluntary running was associated with changes in gut microbial composition. Shot gun metagenomics (MG) was initially performed on 10 sedentary and 10 running individuals from Cohort 1 (**Table S5**), and the relative abundances of the 25 most abundant microbial taxa were compared at genus level (**Figure 4A**). Unclassified genera of the *Lachnospiraceae* family were the only taxonomic bin that differed significantly between running and sedentary mice (**Figure 4B**). Interestingly, *Lachnospiraceae* was also significantly enriched in sedentary HF-fed cohorts in both MG and MT datasets from our previous study of an aging BXD mouse colony (Zhou et al., 2026) (**Figure 4C**).

**Figure 4.**
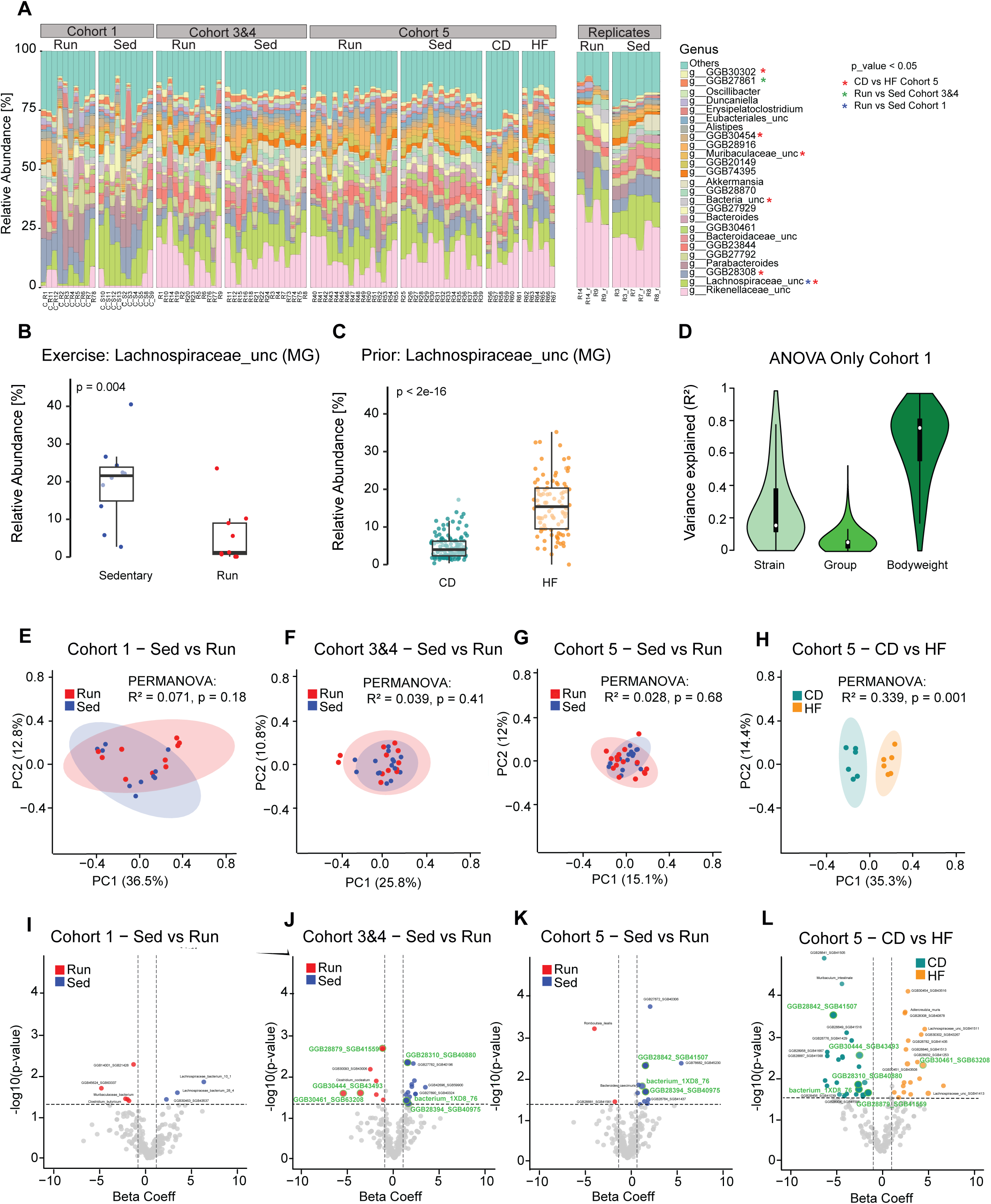
Cecal Microbiome Analysis (**A**) Relative abundances of the 25 most abundant genera in the cecal microbiome across Cohorts 1–5 and the CD versus HF pilot experiment. Significant differences (p < 0.05) between Sed and Run mice are indicated for each cohort, with Cohorts 3 and 4 combined as biological replicates. Significant differences between CD and HF mice in the pilot experiment are also indicated. (**B**) Relative abundance of the *Lachnospiraceae* family, showing a significant decrease in running individuals (p = 0.004). (**C**) This finding is consistent with a recent study of ours (Zhou et al., 2026) in the metagenome (MG) cecal samples of hundreds of mice recorded on either CD or the same HF as used in this study, where HF individuals had a significant increase in *Lachnospiraceae* (p < 2e-16). **(D)** Variance partitioning analysis of species-level abundances in Cohort 1, showing the proportion of variance explained by strain, experimental group, and body-weight AUC. **(E–G)** Principal coordinate analysis (PCoA) of Bray–Curtis dissimilarities at the species level comparing Sed and Run mice in Cohort 1, combined Cohorts 3 and 4, and Cohort 5, respectively. **(H)** PCoA of Bray–Curtis dissimilarities comparing CD- and HF-fed mice in the Cohort 5 pilot experiment.**(I–K)** MaAsLin3 volcano plots showing differentially abundant species between Sed and Run mice in Cohort 1, combined Cohorts 3 and 4, and Cohort 5, respectively. Taxa corresponding to p < 0.05 and |β| ≥ 1 consdered as differentially abundant. **(L)** MaAsLin3 volcano plot showing differentially abundant species between CD- and HF-fed mice in the Cohort 5 pilot experiment.

**Figure S4.**
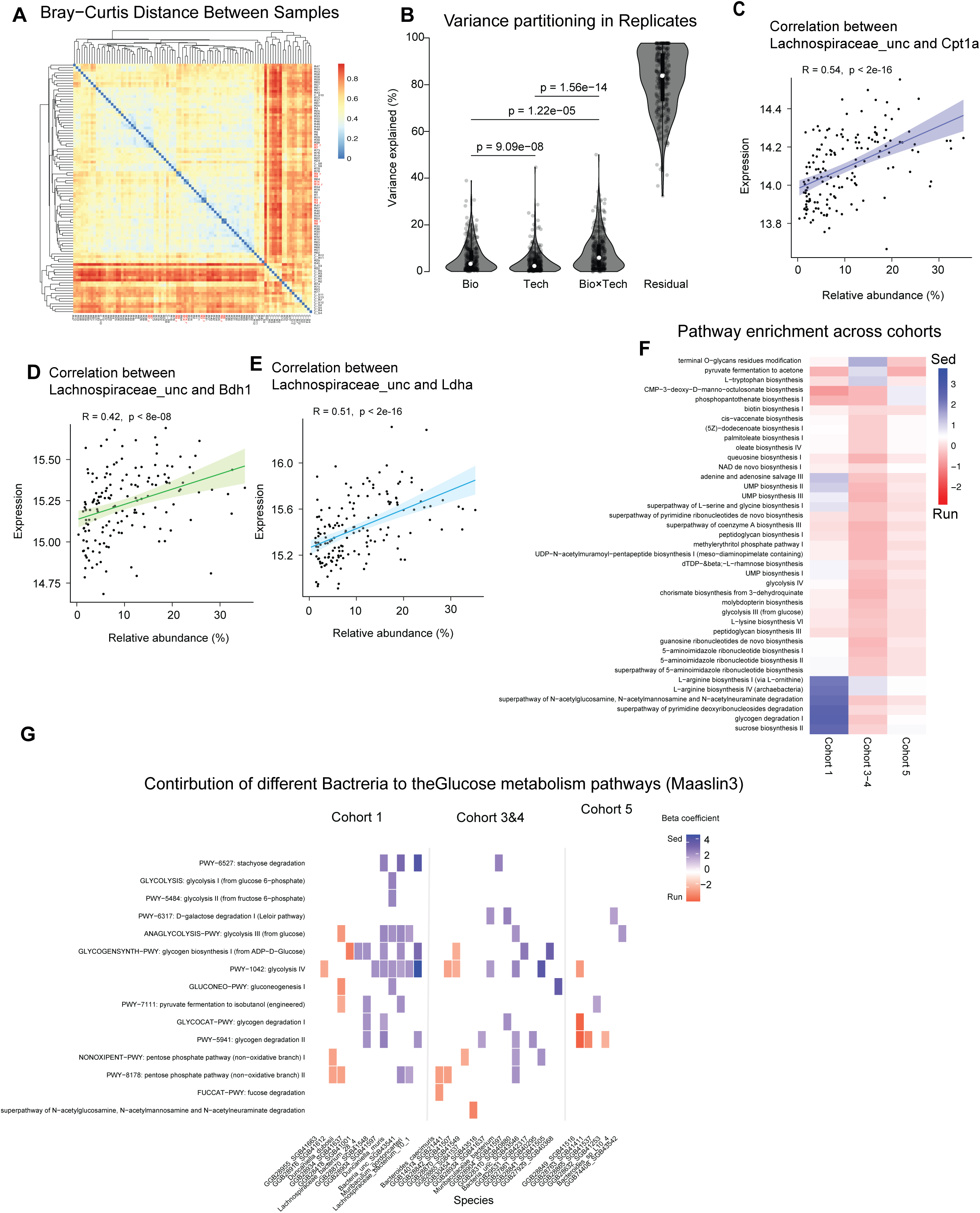
Additional Cecal Microbiome Analysis **(A**) Heatmap of pairwise Bray–Curtis dissimilarities between cecal metagenomic samples, with hierarchical clustering used to assess similarity among biological and technical replicates. (**B**) Variance partitioning of species-level abundances among biological replicates, technical replicates, their interaction, and randomly selected samples. Biological and technical replicates explained significantly less variance than randomly selected samples. (**C–E**) Positive correlations between the relative abundance of *Lachnospiraceae_unclassified* and host cecal expression of (**C**) *Cpt1a* (R = 0.54, p < 2e-16), (**D**) *Bdh1* (R = 0.42, p < 8e-8), and (**E**) *Ldha* (R = 0.51, p < 2e-16). (**F**) Heatmap of differential microbial pathway abundance between Sed and Run mice across Cohort 1, combined Cohorts 3 and 4, and Cohort 5. (**G**) Species-stratified analysis of glucose metabolism-related pathways showing the contribution of individual microbial species to differential pathway abundance between Sed and Run mice across cohorts. Beta coefficients were derived from MaAsLin3 analyses, with positive coefficients indicating enrichment in Sed and negative coefficients indicating enrichment in Run mice.

Using our previously generated microbiome datasets together with host gene-expression data, we further investigated potential associations between *Lachnospiraceae* and host metabolism. *Lachnospiraceae* abundance was significantly correlated with the expression of *Cpt1a* (R= 0.54, p< 2e-16, **Figure S4C**), *Bdh1* (R= 0.42, p< 8e-8, **Figure S4D**), and *Ldha* (R= 0.51, p< 2e-16, **Figure S4E**). These genes are involved in fatty acid oxidation, ketone body metabolism, and glucose/lactate metabolism respectively (Choi et al., 2024; Flack et al., 2018; Williams et al., 2024), and have previously been linked to glucose homeostasis, insulin resistance, and diabetes-related metabolic dysfunction (Lai et al., 2020; Sanchez et al., 2021; Stagg et al., 2021)

To determine whether these exercise-associated microbial differences were reproducible, additional metagenomic sequencing was performed using only BXD43 mice from Cohorts 3 and 4 (**Table S5**). As samples from the different cohorts were processed in separate extraction and sequencing batches (**Table S2**), the datasets were analyzed independently rather than applying computational batch correction across cohorts. Although batch effects can influence the outcome of our analysis, their correction in metagenomic abundance data is challenging because these data are compositional and sparse (Wang & LeCao, 2020). We therefore assessed technical reproducibility directly by including repeated samples across extraction and sequencing batches (**Figure 4A**, **Figure S4A–B**). These repeats showed comparable microbial profiles between runs (**Figure 4A**), with very similar Bray–Curtis distances (**Figure S4A**). ANOVA results further showed that the variance explained by technical replicates was significantly lower than that explained by biological replicates and random samples (p = 1.1e-5 and p = 3e-11, respectively; **Figure S4B**). This supports the evaluation of exercise effects on cecal microbiome composition separately within each cohort, without batch correction.

The exercise-associated difference in *Lachnospiraceae* abundance observed in Cohort 1 was not reproduced in the second sequencing batch comprising BXD43 mice from Cohorts 3&4. Instead, *GGB27861*, a currently uncharacterized genus-level taxonomic bin within Bacteroidetes (Blanco-Miguez et al., 2023), was significantly differentially abundant between running and sedentary BXD43 mice.

PCoA at the species level showed substantial overlap between sedentary and running mice across the independently analyzed cohorts (**Figure 4E–G**). Cohort 1, which included three BXD strains, showed a more dispersed distribution of samples, with some separation between running and sedentary animals but considerable overlap between the two groups and substantial variation within each group (**Figure 4E**). In contrast, samples from Cohorts 3&4 clustered more closely together, with less variation both within and between exercise conditions (**Figure 4F**). MaAsLin3 differential abundance analysis identified a limited number of exercise-associated taxa in Cohort 1 and a larger number in Cohorts 3&4 (**Figure 4I–J**). However, none of the significantly differentially abundant species identified in Cohort 1 were replicated in Cohorts 3&4, indicating that the species-level taxonomic response to exercise was strongly cohort-dependent. This lack of replication may partly reflect differences in the genetic composition of the cohorts, as Cohort 1 comprised three genetically distinct BXD strains, whereas Cohorts 3&4 contained only BXD43 mice. Indeed, within Cohort 1, genetic background explained approximately 15% of the variation in microbial composition, compared with approximately 5% explained by exercise (**Figure 4D**). The greater influence of genetic background may also contribute to the more dispersed distribution of samples observed in the PCoA for Cohort 1(**Figure 4E**).

We next examined BXD43 mice from Cohort 5, in which approximately 0.5 L of soiled bedding was reciprocally exchanged weekly between paired sedentary and running cages from week 10 onwards (**Table S5**). Sedentary and running animals showed particularly tight clustering and reduced dispersion by PCoA compared with Cohort 1 and Cohorts 3&4 (**Figure 4G**). Consistently, MaAsLin3 identified fewer differentially abundant taxa between sedentary and running mice in Cohort 5 compared to Cohort 3&4. At the species level, only two exercise-associated taxa were shared between Cohorts 3&4 and Cohort 5, namely *bacterium-1XD8_76* and *GGB28394_SGB40975* (**Figure 4K**).

We also performed functional profiling of the cecal microbiome from Cohort 1, Cohorts 3&4, and Cohort 5 using HUMAnN3 (**Table S6**), followed by MaAsLin3 analysis. Several MetaCyc pathways enriched in sedentary mice in Cohort 1, including glycogen degradation and sucrose biosynthesis, showed the opposite association in Cohorts 3&4, with grater abundance in running mice, potentially reflecting differences in genetic background (**Figure S4F**). Interestingly, Cohorts 3&4 and Cohort 5, which differed primarily by the bedding-exchange intervention, showed broadly similar pathway associations, although effect sizes were generally reduced in Cohort 5.

Analysis of the contribution of individual bacterial species to specific glucose-related pathways revealed considerable variation across cohorts (**Figure S4G**). Cohorts 3&4 showed contributions from multiple species across a broad range of pathways, whereas fewer species–pathway associations were observed in Cohort 5. Nevertheless, several pathways remained represented in both cohorts, including glycolysis and glycogen degradation, although the contributing species and direction of their associations differed. This reduction in species-specific associations in Cohort 5 is consistent with an attenuation of exercise-associated microbial differences following bedding exchange. As a relaxed q-value cutoff of **q < 0.25** was used for this exploratory analysis, these results should be interpreted as trends rather than statistically significant associations. However, three pathways, terminal O-glycan residue modification, pyruvate fermentation to acetone, and L-tryptophan biosynthesis, showed opposite associations, being enriched in sedentary mice in Cohorts 3&4 but in running mice in Cohort 5.

A small parallel experiment using sedentary BXD43 mice maintained on CD or HF was performed to validate the effectiveness of the bedding-exchange procedure. As our previous study showed pronounced diet-associated differences in gut microbiome composition between CD- and HF-fed mice (Zhou et al., 2026), these animals provided a model in which microbial transfer between cages could be more readily assessed. Despite the uncontrolled microbiome transfer introduced by bedding exchange, CD- and HF-fed mice retained a pronounced separation by PCoA (**Figure 4H**) and showed a larger number of differentially abundant taxa by MaAsLin3 compared with the Sed–Run comparison (**Figure 4L**). Some taxa identified in the Sed–Run comparisons were also differentially abundant between CD and HF animals, although the overall diet-associated microbial differences were substantially greater than those associated with exercise.

### Longitudinal dynamics of the fecal microbiome during the bedding exchange experiment

Longitudinal fecal metagenomic sequencing at weeks 8, 9, 11, 13, 16, 19, and 31 allowed us to follow microbiome dynamics throughout the bedding-exchange experiment (**Figure S5A, Table S7 and S8**).

Simpson diversity was initially comparable across groups but decreased following the transition to HF feeding, while remaining higher in the CD group (**Figure S5B**). At later time points, these differences became less pronounced, with no significant differences between Tablegroups observed by week 31.

At the genus level, microbial composition showed considerable variation over time, but no clear separation between the HF, running, and sedentary groups was apparent at individual time points (**Figure S5C**). In contrast, the CD group maintained a more distinct microbial composition, particularly during the earlier time points.

Interestingly, individual bacterial taxa showed distinct patterns over the course of the experiment (**Figure S5D**). For example, *GGB28632* showed more similar abundance trajectories between groups undergoing reciprocal bedding exchange, suggesting a potential influence of microbial transfer. In contrast, *GGB28308* showed more similar trajectories between groups maintained on the same diet, suggesting that its abundance was more strongly associated with dietary conditions than with bedding exchange. As bedding exchange represents an uncontrolled form of microbial transfer, its efficiency may differ between taxa, particularly depending on their ability to survive exposure to oxygen during bedding collection and transfer, potentially favoring more aerotolerant bacteria over strict anaerobes(Gheorghe et al., 2021).

Consistent with this temporal variability, we next quantified the proportion of overall microbial variation associated with exercise at each fecal time point and in the endpoint, cecum using Bray–Curtis PERMANOVA (**Figure S5E**). Exercise-associated variance in the fecal microbiome varied over time, increasing from approximately 5–10% during the earlier time points to approximately 13–14% at weeks 9–12 before decreasing again by week 31. Individual Sed–Run PERMANOVA comparisons were generally not statistically significant, with week 19 being the only significant time point (p = 0.01), and these variance estimates were therefore considered exploratory. In contrast, exercise explained only approximately 2% of the variation in the endpoint cecal microbiome.

To further characterize the temporal dynamics underlying these patterns, we performed longitudinal analysis within the running and sedentary groups using MaAsLin3. The most pronounced shift in microbial composition occurred between week 8 and the following week, coinciding with the introduction of the HF diet (**Figure S5F**). Numerous taxa changed in abundance within this single one-week interval, demonstrating that HF feeding can rapidly alter the gut microbiome. In contrast, comparisons between subsequent consecutive time points showed fewer and less consistent changes, with individual taxa increasing or decreasing at different stages of the experiment MaAsLin3 analysis of sedentary and running mice at each time point revealed a time-dependent effect of exercise on microbial composition (**Figure S5G**). Relatively few differentially abundant taxa were detected at earlier time points, whereas the largest number of Sed–Run differences was observed at week 19, with few differences also present at week 31. The taxa identified varied considerably between time points, indicating that exercise-associated differences in the fecal microbiome were dynamic rather than consistently maintained throughout the experiment.

Overall, the longitudinal analysis indicated a greater and temporally variable exercise-associated component in the fecal microbiome than in the endpoint cecal microbiome.

### Bedding-Exchange Cohort: Testing Microbiome Causality

The bedding Exchange Cohort allowed us to test whether the microbial alterations associated with voluntary physical activity were sufficient to influence metabolic outcomes independently of exercise behavior itself, or whether they represented a downstream consequence of the exercise-diet interaction rather than a causal mediator.

As in the previous cohorts, voluntary running activity progressively declined with age in Cohort 5 (p = 1e-3, **Figure 5A**). Median weekly running distance decreased from approximately 57 km/week at 8–18 weeks of age to 38 km/week at 19–29 weeks (p = 3e-3; **Figure 5B**). Overall wheel activity in Cohort 5 was comparable to that previously observed in BXD43 mice from Cohorts 1–4 (p = 0.2; **Figure 5C)** and remained substantially higher than in the other BXD strains (p = 1e-7; **Figure 5C**). Home-cage movement similarly decreased with age (p = 8e-6; **Figure 5D**). Median weekly locomotion significantly dropped between week 14-20 and weeks 23-29 (p=2e-5; **Figure 5E**). Total home-cage locomotion was significantly lower in Cohort 5 compared to BXD43s of Cohorts 1-4, while overall home-cage activity in Cohort 5 did not significantly differ from that of the other strains of Cohorts 1–4 (p = 5e-4, p = 0.2; **Figure 5F**).

**Figure 5.**
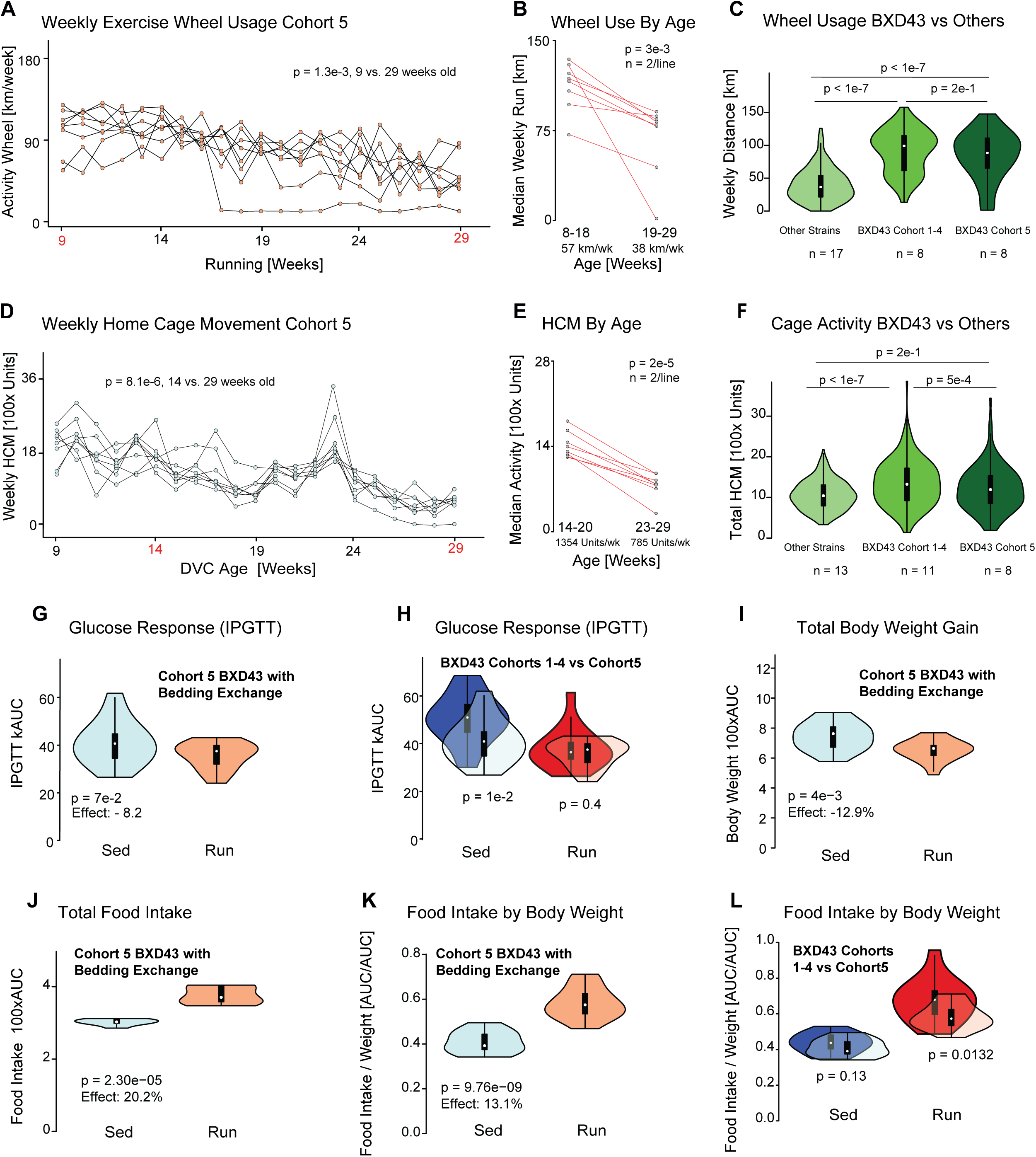
Physical Activity and Metabolic Phenotypes of Cohort 5 with Bedding Exchange (**A**) Weekly running-wheel activity in Cohort 5, showing a significant decline with age (p = 1.3e-3). (**B**) Comparison of median weekly running distance between weeks 8–18 and 19– 29, showing a significant decrease from 57 to 38 km/week (p = 3e-3). (**C**) Weekly running distance in BXD43 mice from Cohort 5 compared with BXD43 mice from Cohorts 1–4 and the other strains, confirming the higher running activity of BXD43 mice. (**D**) Weekly home-cage monitoring (HCM) activity in Cohort 5, showing a significant decline with age (p = 8.1e-6). (**E**) Median HCM activity decreased from 1,354 to 785 units/week between weeks 14–20 and 23–29 (p = 2e-5). (**F**) Comparison of total HCM activity between BXD43 mice from Cohort 5, BXD43 mice from Cohorts 1–4, and the other strains. (**G**) IPGTT glucose AUC was 8.2% lower in Run compared with Sed mice in Cohort 5, although the difference did not reach statistical significance (p = 7e-2). (**H**) Comparison of IPGTT glucose AUC between Cohort 5 and BXD43 mice from Cohorts 1–4, showing a significant difference between sedentary groups (p = 1e-2), but not between running groups (p = 0.4). (**I**) Body-weight AUC was 12.9% lower in Run compared with Sed mice (p = 4e-3). (**J**) Total food-intake AUC was 20.2% higher in Run mice (p = 2.30e-5). (**K**) Food intake-to-body-weight AUC ratio was 13.1% higher in Run mice (p = 9.76e-9). (**L**) Comparison of the food intake-to-body-weight AUC ratio between Cohort 5 and BXD43 mice from Cohorts 1–4, showing no significant difference between sedentary groups (p = 0.13) but a significant difference between running groups (p = 0.0132).

**Figure S5.**
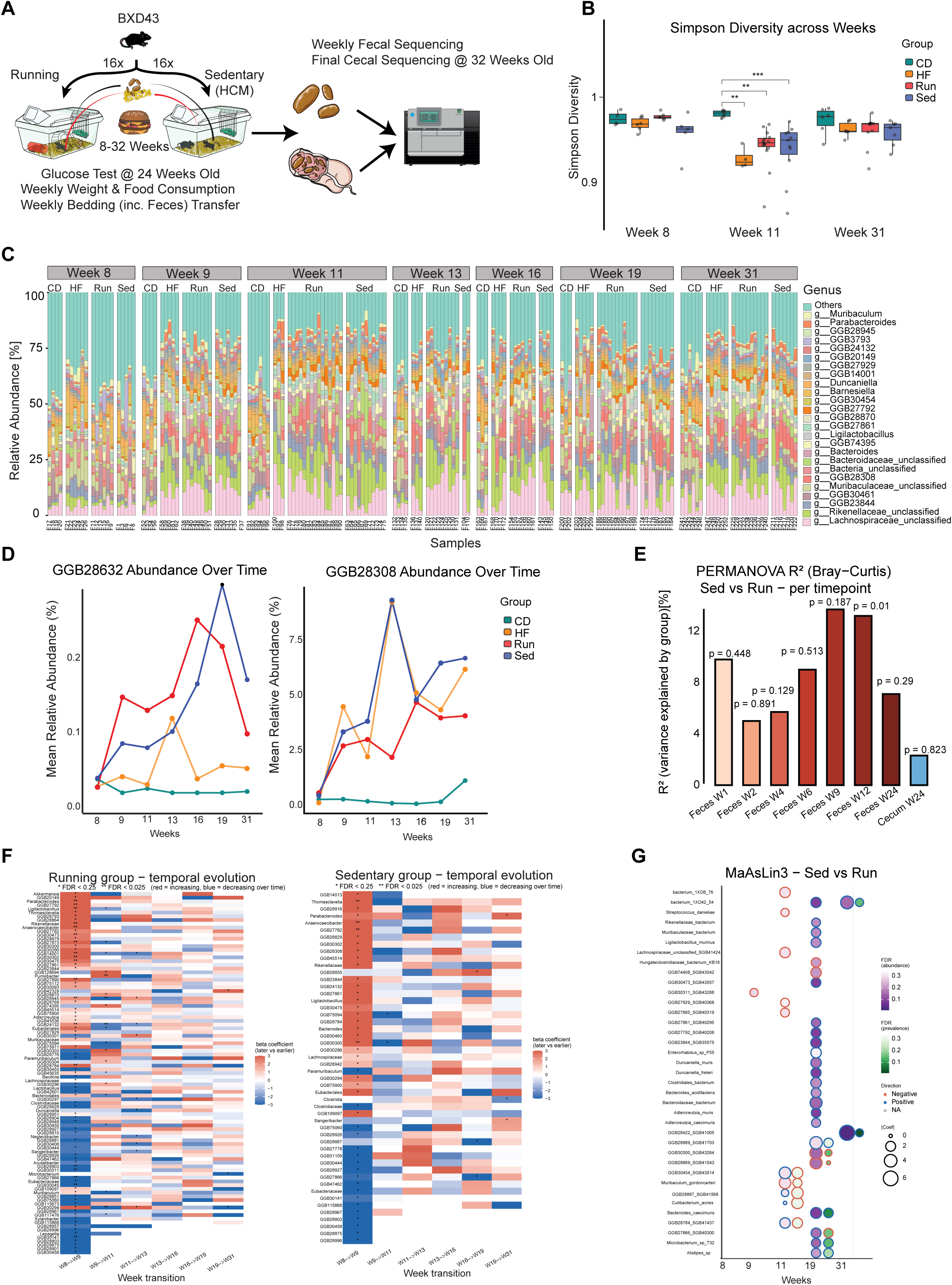
Longitudinal Fecal Microbiome Analysis in Cohort 5 (**A**) Study design of an additional cohort (Cohort 5) consisting exclusively of BXD43 mice, differing from the previous cohorts by the implementation of weekly reciprocal bedding exchange and longitudinal fecal collection for microbiome composition analysis. (**B**) Simpson diversity across weeks 8, 11, and 31, showing changes in within-sample microbial diversity over time (**C**) Relative abundances of the most abundant genera in fecal samples from CD, HF, Sed, and Run mice across the longitudinal sampling timepoints. (**D**) Longitudinal relative abundance of selected bacterial taxa across CD, HF, Sed, and Run groups. (**E**) Proportion of microbiome compositional variation explained by exercise at each fecal timepoint and in the endpoint cecum, determined by Bray–Curtis PERMANOVA. Exercise explained approximately 5–14% of fecal microbiome variation across time but only ∼2% in the endpoint cecum; individual Sed–Run comparisons were not significant except at week 19 (p = 0.01). (**F**) MaAsLin3 analysis of temporal changes in species abundance within Run and Sed mice separately. Consecutive sampling timepoints were compared, and species with an FDR-adjusted q-value < 0.25 in at least one comparison are shown; red and blue indicate increasing and decreasing abundance over time, respectively. (**G**) MaAsLin3 analysis comparing Sed and Run mice at individual fecal sampling timepoints. Species reaching a joint FDR-adjusted q-value < 0.35 at at least one timepoint are shown, with significant associations observed only at week 19.

We next examined whether the metabolic phenotype associated with voluntary running was maintained following reciprocal bedding exchange. Within Cohort 5, running mice showed an 8.2% lower glucose AUC than sedentary mice, although this difference did not reach statistical significance (p = 0.07; **Figure 5G**). Comparison with BXD43 mice from Cohorts 1– 4 revealed a marked change in the glucose-response phenotype (**Figure 5H**). Sedentary mice in Cohort 5 showed an improved glucose response compared with sedentary BXD43 mice from Cohorts 1–4 (p = 1e-2). Consequently, the pronounced difference in glucose response between sedentary and running BXD43 mice observed in Cohorts 1–4 was strongly reduced in Cohort 5. Running animals nevertheless continued to show differences in body-weight gain and food intake compared with sedentary animals, although these effects were attenuated relative to BXD43 mice from Cohorts 1–4. Running mice gained significantly less body weight than sedentary animals over the experimental period, with a 12.9% reduction in body-weight AUC (*p* = 0.004; **Figure 5I**). This difference was approximately half of that observed in BXD43 mice from Cohort 4, in which running was associated with an approximately 25% reduction in body-weight AUC (**Figure S2G**). Despite their lower weight gain, Cohort 5 running mice consumed 20.2% more food than sedentary animals (*p* = 2e-5; **Figure 5J**), corresponding to a 13.1% higher food intake-to-body-weight ratio (*p* = 1e-8; **Figure 5K**). Comparison with BXD43 mice from Cohorts 1–4 showed that the food intake to body weight ratio was comparable between sedentary groups (*p* = 0.13, **Figure 5L**), whereas it was significantly lower in Cohort 5 running mice than in running mice from Cohorts 1–4 (*p* = 0.01; **Figure 5L**). Thus, although running mice in Cohort 5 maintained lower weight gain and higher food intake than their sedentary counterparts, both the reduction in weight gain and the exercise-associated increase in food intake relative to body weight were attenuated compared with the BXD43s from previous cohorts.

Overall, the effects of running on body-weight gain, food intake, and particularly glucose response were attenuated in Cohort 5 compared with BXD43 mice from Cohorts 1–4, coinciding with our hypothesis regarding the reciprocal bedding exchange.

### Circadian and Procedure-Associated Patterns of Home-Cage Locomotion and Running-Wheel Activity

Home-cage activity was recorded using an electrode-based monitoring system positioned underneath the DVC cages and was consistently higher during the dark phase (7 pm–7 am) than during the light phase (7 am–7 pm). This pattern is clearly visible across consecutive days in the activity heatmap (example day 49-54, **Figure 6A**). Quantification of home cage activity across all cohorts confirmed this difference, with median locomotor activity nearly tripling during the night (*p* < 1e-4; **Figure 6B**). Running-wheel activity followed a similar day–night pattern, with most running occurring during the dark phase (example day 40-54; **Figure 7A**). Accordingly, mice run significantly greater distances during the night than during the day (*p* = 8e-3; **Figure 7B**).

**Figure 6.**
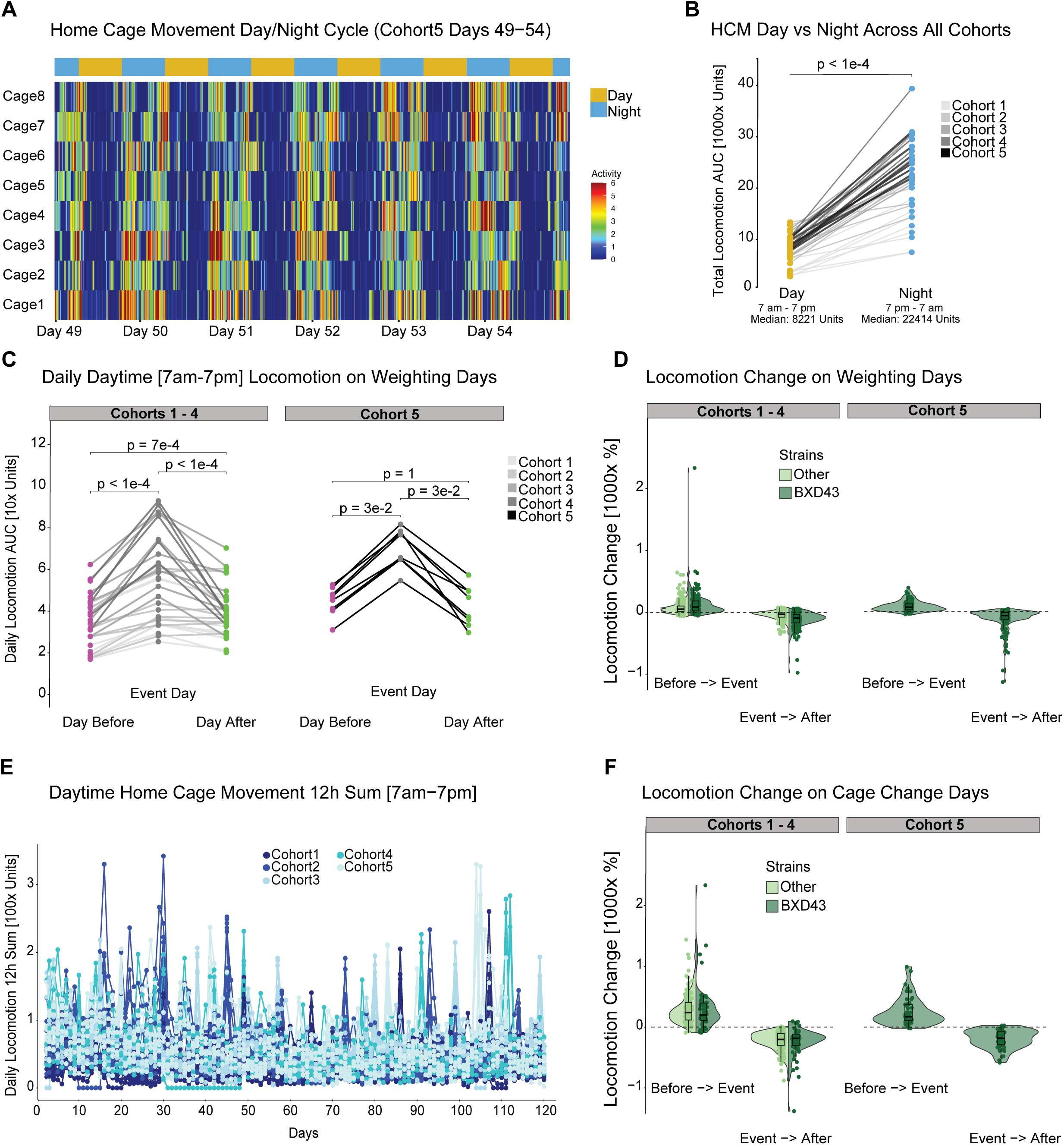
Home Cage Locomotion Analysis (**A**) Representative home-cage monitoring (HCM) activity profiles across the day–night cycle in Cohort 5, illustrating the higher locomotor activity during the dark phase. (**B**) Comparison of daytime (7 am–7 pm) and nighttime (7 pm–7 am) HCM activity across all cohorts. Locomotor activity was significantly higher during the night, with median values of 22,414 units at night and 8,221 units during the day (p < 1e-4). (**C**) Daytime locomotor activity on the day before, day of, and day after weighing across Cohorts 1–5, showing changes in activity associated with the weighing procedure. (**D**) Relative change in locomotor activity from the day before to the weighing day and from the weighing day to the following day, comparing BXD43 mice with the other strains and Cohort 5 with Cohorts 1–4. (**E**) Daily locomotor activity around cage-change days in Cohorts 1–4 and Cohort 5. (**F**) Relative change in locomotor activity from the day before to the cage-change day and from the cage-change day to the following day, comparing BXD43 mice with the other strains and Cohort 5 with Cohorts 1–4. P-values for the corresponding comparisons are indicated in the panels.

**Figure 7.**
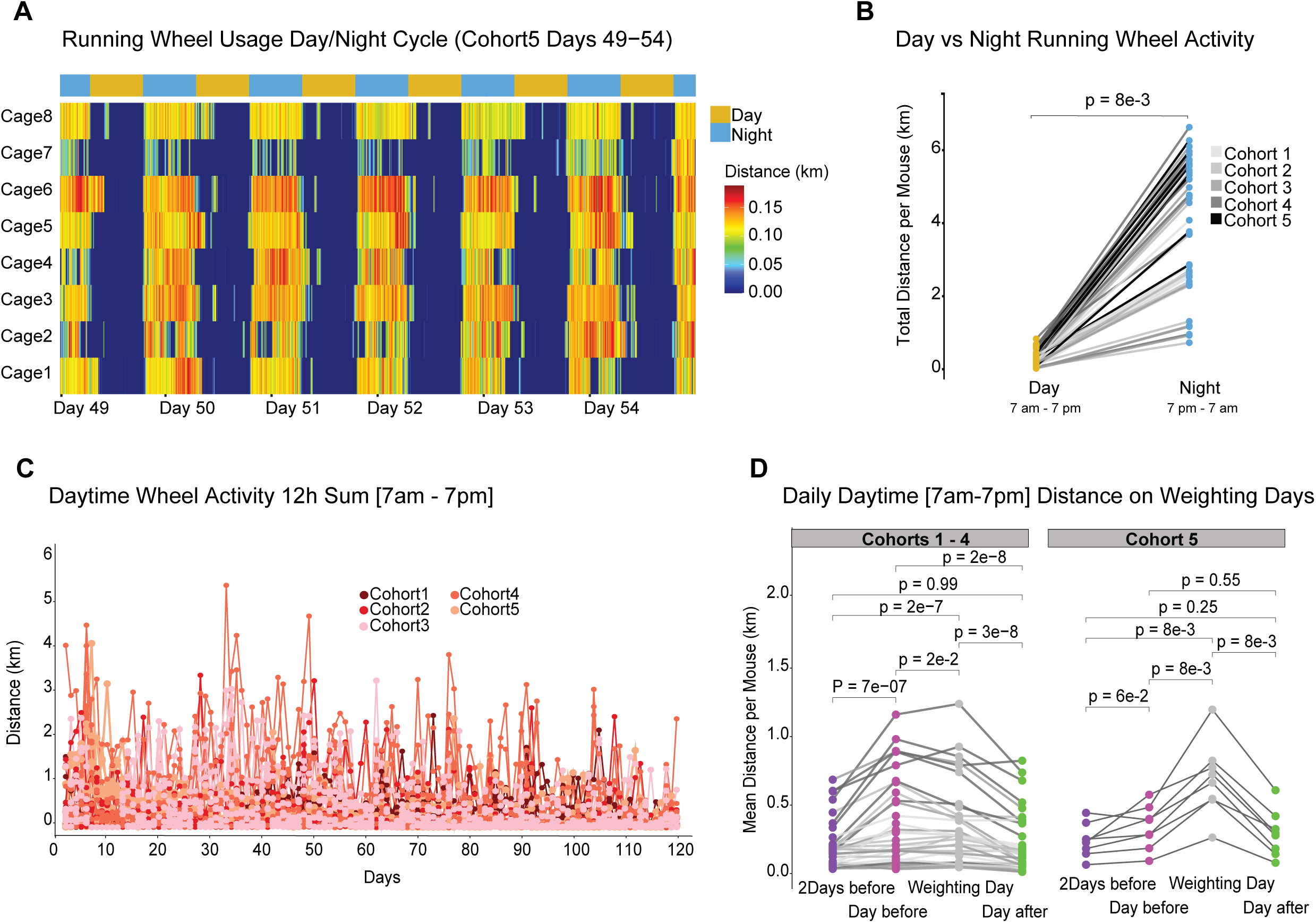
Running Wheel Usage (**A**) Representative running-wheel activity across the day–night cycle in Cohort 5 (days 49– 54), showing the predominantly nocturnal pattern of voluntary running. (**B**) Comparison of running distance during the day (7 am–7 pm) and night (7 pm–7 am) across all running cohorts, showing significantly greater wheel activity during the night (p = 8e-3). (**C**) Daily daytime running distance (7 am–7 pm) across the experimental period for all running cohorts with higher peaks corresponding to the cage change days. (**D**) Daytime running distance around weighing days in Cohorts 1–4 and Cohort 5. In Cohorts 1–4, running activity increased from two days before to the day before weighing (p = 7e-7), increased slightly further on the weighing day (p = 2e-2), and decreased the following day (p = 3e-8). In Cohort 5, no significant increase was observed on the day before weighing (p = 6e-2), whereas activity increased on the weighing day (p = 8e-3) and decreased the following day (p = 8e-3).

We next examined whether routine experimental procedures were associated with changes in daytime activity. In Cohorts 1–4, daytime home-cage locomotion increased significantly on weighing days compared with both the preceding and following days (both p < 1e-4; **Figure 6C**). A similar pattern was observed in Cohort 5, where locomotor activity increased from the day before to the weighing day (p = 0.03; **Figure 6C**) and decreased again on the following day (p = 0.03; **Figure 6C**). In Cohort 5, weighing coincided with reciprocal bedding transfer and fecal collection, during which mice were temporarily isolated to make sure that fecal samples could be assigned to the correct individual. Despite these additional perturbations, the transient change in locomotor activity was comparable between Cohort 5 and Cohorts 1–4 (**Figure 6D**). Days coinciding with routine DVC cage changes, which were performed every three weeks, were excluded from the weighing-day analysis and analyzed separately.

Across the experimental period, temporal daytime home-cage locomotion curves showed several temporary increases in activity, with the largest peaks occurring on cage-change days (**Figure 6E**). In fact, there is an increase in activity from the preceding day to the cage-change day, followed by a decrease on the subsequent day in both Cohorts 1–4 and Cohort 5 (**Figure 6F**). Although a similar transient response was observed around the weighing days (**Figure 6D**), cage changes appeared to have a greater impact on daytime home-cage locomotion. This is also true for Cohort 5, suggesting that these additional procedures had a smaller impact on daytime home-cage activity.

Running-wheel activity showed similar transient changes in response to experimental procedures. Also, temporal daytime wheel-running activity curves showed recurrent peaks that coincided with scheduled cage-change days (**Figure 7C**), similar to the pattern observed for home-cage locomotion (**Figure 6E**). In Cohorts 1–4, daytime running increased markedly from two days before to the day preceding weighing (p = 7e-7; **Figure 7D**), coinciding with the routine cage change performed by animal-care staff. Running remained elevated on the weighing day, before decreasing again on the following day (p = 3e-8; **Figure 7D**). In Cohort 5, cage changes were instead performed on the weighing day, together with reciprocal bedding transfer, and feces collection. No significant change in running was observed between two days before and the day before (p = 6e-2; **Figure 7D**). Instead, running increased on the weighing day (*p* = 8e-3) and decreased again on the following day (*p* = 8e-3; **Figure 7D**).

Overall, both home-cage locomotion and voluntary wheel running were predominantly nocturnal and showed transient increases in response to routine experimental procedures. Cage changes had the greatest impact on activity, whereas the additional bedding exchange and fecal collection procedures in Cohort 5 had a comparatively smaller effect.

## DISCUSSION

Gene–environment interactions are known to play a major role in nearly all chronic non-communicable diseases (Herrera-Luis et al., 2024). In this study, we identified an inbred mouse strain, BXD43, which is exceptionally active in running and is able to substantially, although not entirely mitigate the effects of HF feeding on obesity and metabolic dysfunction. While exercising mice had significantly increased food intake compared with sedentary mice, their metabolic health was also significantly improved, despite relatively little effect on body weight.

This is particularly interesting considering the large variability in the response to lifestyle interventions observed in humans. Low-calorie diets and physical activity are commonly used to prevent and treat obesity and related metabolic disorders such as type 2 diabetes, but their effectiveness varies considerably between individuals. Some people manage to lose substantial amounts of weight, whereas others show only limited weight loss despite similar interventions and efforts (Dent et al., 2020; Yaskolka Meir et al., 2025). Several intrinsic and extrinsic factors may contribute to this variability, including genetic background, metabolic differences, and environmental factors (Dent et al., 2020). The three BXD strains chosen for our study also showed distinct responses to HF feeding. BXD100 was relatively heavy independently of diet, whereas BXD43 showed a pronounced increase in body weight in response to HF feeding. In contrast, BXD40 remained relatively lean even under HF feeding.

The substantial differences in voluntary running between the BXD strains further highlight the influence of genetic background on exercise behavior (Avila et al., 2017; Lightfoot et al., 2004). We therefore focused on BXD43, the most active strain, to investigate whether its exceptionally high level of voluntary exercise would result in stronger metabolic effects. BXD43 maintained considerably higher voluntary running activity than the other BXD strains, even after wheel usage decreased with age. Interestingly, this high voluntary running activity did not correspond to similarly increased spontaneous home-cage locomotion. This suggests that voluntary wheel running and normal home-cage movement are at least partly distinct behavioral traits and that high spontaneous locomotion does not necessarily predict how much an animal will voluntarily exercise when given access to a wheel, which is in concordance with a previous study (Niiranen et al., 2023). The progressive decline in wheel running over the course of the experiment is also consistent with previous studies showing that voluntary wheel use decreases with age and duration of wheel exposure (Figueiredo et al., 2009; Jung & Luthin, 2010). Nevertheless, BXD43 mice continued to run substantial distances throughout the experiment, making this strain particularly suitable for studying the long-term effects of voluntary exercise.

One of the most striking findings was that these high levels of voluntary exercise substantially improved glucose response despite relatively modest effects on body weight. Running mice also consumed more food than sedentary mice, with the difference becoming even larger when looking at the food intake to bodyweight ratio. Increased food intake may therefore partly compensate for the additional energy expenditure caused by exercise and explain why the large amount of physical activity did not result in similarly large reductions in body weight. This is consistent with previous observations that voluntary exercise can increase energy intake in mice (Kelly et al., 2011). Importantly, the improved glucose response was maintained despite this increased caloric intake, indicating that the metabolic benefits of exercise in our mice were not simply a consequence of weight loss, but more due to metabolic plasticity.

This separation between body weight and glucose response is particularly relevant because exercise can influence glucose homeostasis through several mechanisms independent of changes in body weight. Exercise increases glucose uptake by skeletal muscle and improves insulin sensitivity (Sylow et al., 2017), while other mechanisms, including hypothalamic regulation and changes in inflammatory signaling, can also influence glucose homeostasis independently of adiposity (Douglass et al., 2023; Sasaki et al., 2024). This may also help explain the unexpected inverse association between body weight and glucose response observed in both sedentary and running mice. Although higher body weight and adiposity are generally associated with impaired glucose homeostasis, these traits are not always directly coupled.

We next investigated whether the gut microbiome could contribute to some of these exercise-associated metabolic effects. In contrast to the relatively pronounced metabolic phenotype, exercise produced only modest and cohort-dependent changes in cecal microbial composition. Previous studies have shown that voluntary exercise can alter the mouse gut microbiome, including under HF feeding (Allen et al., 2015; Evans et al., 2014).

However, in our study we could not identify a specific taxon that was consistently differentially abundant with exercise across cohorts. In Cohort 1, unclassified taxonomic bins belonging to the *Lachnospiraceae* family were reduced in running compared with sedentary mice, but this difference was not reproduced in the other cohorts. Interestingly, members of *Lachnospiraceae* have repeatedly been associated with exercise, although the direction of the response differs between studies. Evans et al. (2014) found that voluntary wheel running increased *Lachnospiraceae* abundance, whereas in our Cohort 1 the unclassified Lachnospiraceae-related taxon was reduced in running mice. Similarly, Zhang et al. (2026) reported an increase in an unclassified *Lachnospiraceae* genus following exercise. Lamoureux et al. (2017) also identified *Lachnospiraceae* among the taxa associated with exercise using a machine-learning approach, although conventional differential abundance analyses did not detect significant exercise-associated changes. These contrasting findings suggest that the response of *Lachnospiraceae* to exercise is dependent on the experimental context rather than showing a consistent direction across studies. These contrasting findings suggest that although exercise can induce an overall shift in microbial composition, the individual taxa driving this shift varies between studies depending on the experimental context. Thus, there may not be a single taxonomic signature that consistently characterizes the microbial response to exercise. Diet and genetic background may contribute to this variability. In Cohort 1, which contained BXD40, BXD43 and BXD100 mice, genetic background explained approximately 15% of the variation in microbial composition, whereas exercise explained only approximately 5%. This is consistent with previous evidence that host genetics can influence gut microbial composition (McKnite et al., 2012; Perez-Munoz et al., 2019). Importantly, using 43 genetically distinct BXD strains and the same HF diet used in our previous study, we showed that diet was the major determinants of cecal microbial composition, with HF feeding also affecting taxa belonging to *Lachnospiraceae* (Zhou et al., 2026). The stronger effect of diet compared with exercise may reflect its more direct influence on the gut microbial environment, as diet directly determines nutrient and substrate availability and can also alter bile-acid metabolism and intestinal conditions (Daniel et al., 2014).

In comparison, the effect of exercise on the gut microbiome is likely to be more indirect. Exercise can modify host factors that influence the intestinal environment, including intestinal motility and transit time, gastrointestinal blood flow, bile-acid metabolism, immune signaling and hyperthermia (Alsinani et al., 2026). The microbiome may therefore respond secondarily to physiological changes induced by exercise. This could partly explain why exercise-associated taxonomic changes tend to be relatively modest and variable between studies, and why we could not identify a specific taxon that was consistently altered across our independent cohorts.

The intestinal sampling site may represent an additional source of variability. Exercise-associated microbial changes have been shown to be gut-segment dependent, with differences in both the magnitude of the response and the taxa affected across intestinal regions (Denou et al., 2016). This may partly explain the relatively small exercise-associated effect observed in our endpoint cecal samples and highlights the importance of examining longitudinal fecal samples.

The longitudinal fecal microbiome analysis further showed that the microbial response was dynamic rather than constant throughout the experiment. The strongest overall microbial changes occurred shortly after the introduction of the HF diet, demonstrating how rapid diet can modify the gut microbiome. Exercise-associated variation also changed over time, reaching a maximum of approximately 13–14% at week 19 before decreasing later at the end of the study. However, these estimates should be considered exploratory, as most individual Sed–Run PERMANOVA comparisons were not statistically significant. Interestingly, running distance also progressively decreased during the experiment, raising the possibility that the magnitude of exercise-associated microbial differences may change with running activity. Exercise explained only approximately 2% of endpoint cecal variation, further suggesting that exercise-associated effects may differ between fecal and cecal communities.

Despite the limited taxonomic differences, functional profiling identified exercise-associated differences in several MetaCyc microbial pathways involved in carbohydrate and lipid metabolism, and in nucleotide and amino-acids biosynthesis. The direction of some of these associations and the bacterial species contributing to individual pathways differed between cohorts. This suggests that similar microbial functions may be performed by different bacterial communities and that functional changes may therefore be more informative than individual exercise-associated taxa. However, these analyses were exploratory and should be interpreted cautiously. Consistent with a possible functional contribution of the microbiome, exercise-associated remodeling of the gut microbiome and microbial metabolites was observed, and transplantation of microbiota from exercised animals reduced HF-induced obesity and improved glucose homeostasis and insulin sensitivity in recipient mice (Zhang et al., 2026).

The reciprocal bedding-exchange experiment provided an additional way to investigate the potential relationship between the microbiome and the metabolic effects of exercise. Repeated transfer of dirty bedding can promote microbial exchange between mice because mice are naturally coprophagic and are continuously exposed to fecal material present in their environment. Bedding and housing conditions have previously been shown to influence gut microbial composition, and bedding transfer has been used to promote microbial exchange between experimental groups (Miyoshi et al., 2018; Ericsson et al., 2018; Gheorghe et al., 2021). In Cohort 5, sedentary and running BXD43 mice showed tighter microbial clustering and fewer differentially abundant taxa than the independent BXD43 cohorts, consistent with reduced microbial differentiation following weekly bedding exchange.

Interestingly, this reduced microbial differentiation coincided with a marked attenuation of the metabolic differences between sedentary and running BXD43 mice. The pronounced difference in glucose response observed in Cohorts 1–4 was strongly reduced in Cohort 5. Sedentary Cohort 5 mice showed an improved glucose response compared with sedentary BXD43 mice from the previous cohorts. Effects on body-weight gain and food intake were also reduced compared with the previous BXD43s, although running mice still gained less weight and consumed more food than their sedentary counterparts. This parallel reduction in microbial and metabolic differences is consistent with the possibility that microbial exchange contributed to the altered metabolic phenotype.

Running-wheel activity and home-cage locomotion were comparable between Cohort 5 and the previous BXD43 cohorts. Continuous monitoring also showed that both home-cage locomotion and wheel running remained predominantly nocturnal and that routine experimental procedures caused only transient increases in activity. Cage changes produced the largest apparent response, whereas the additional bedding exchange and fecal collection performed in Cohort 5 were associated with smaller changes comparable to those of the other Cohorts. Thus, there was no indication that the bedding-exchange intervention substantially influenced voluntary physical activity or locomotor behavior. Nevertheless, the bedding-exchange experiment has important limitations. Reciprocal soiled bedding exchange represents an uncontrolled form of microbial transfer, and not all bacterial taxa are likely to be transferred and established equally efficiently. Factors such as oxygen sensitivity and the ability to colonize the recipient gut may influence which microorganisms are successfully exchanged (Chu et al., 2017; Gheorghe et al., 2021; Podlesny et al., 2022). However, this variability is not specific to bedding exchange, as even more controlled approaches such as fecal microbiota transplantation by oral gavage do not result in uniform establishment of donor microorganisms, with colonization depending on both donor and recipient characteristics (Walter et al., 2021; Podlesny et al., 2022).

Moreover, soiled bedding contains not only microorganisms but also urine, fecal metabolites, pheromones and other biologically active environmental materials. Urine and feces contain olfactory signals that can be transferred with soiled bedding and potentially influence the behavior and physiology of recipient mice (Merley et al., 2022). The additional handling associated with bedding exchange and fecal collection may also induce transient stress responses and cannot be completely separated from the intervention. Therefore, although reduced microbial differentiation coincides with convergence of the glucose phenotype, contributions from non-microbial components of the bedding or the intervention itself cannot be excluded, and the present experiment cannot establish that the sole microbial transfer caused this metabolic change.

Overall, our findings demonstrate that long-term voluntary exercise can substantially improve glucose metabolism during HF feeding despite increased caloric intake and relatively modest effects on body weight. BXD43 represents a particularly active model for studying the long-term effects of voluntary exercise. Exercise-associated microbiome changes were comparatively subtle, dynamic, and dependent on cohort and sampling site, while functional differences suggest that microbial metabolic activity may be more informative than individual bacterial taxa. The reduced microbial and metabolic differences observed following reciprocal bedding exchange further support a potential contribution of the gut microbiome to the response to exercise, although this relationship remains to be experimentally confirmed. Together, our findings highlight the complex interaction between genetic background, exercise and the gut microbiome in shaping metabolic responses to HF feeding.

## MATERIALS & METHODS

### Animal Handling and Cohort Selection

The 3 strains selected for this study were ordered from The Jackson Laboratory and imported to our facility at the University of Luxembourg in fall 2020. All strains went through one to two rounds of breeding in Specific Pathogen Free (SPF) conditions to amplify the line and create sufficient embryos for embryo transfer to our Specific and Opportunistic Pathogen Free (SOPF) facility for long-term study maintenance. In March 2021, our study protocol, GEDI:CAPMED, was approved by the Luxembourgish Ministry of the Environment (protocol LUPA 2021/07) and the first of five cohorts, all starting with 32 mice, began in May 2022 and concluded in November 2025. The 32 mice of each cohort were separated evenly into those with *ad libitum* access to a recorded running wheel, and those with simple home cage monitoring, without a wheel (**Figure 1D**). All mice were established in cages of 2 individuals/housing unit, which was intended to minimize conflicts over running wheel usage and saturation of home cage monitoring sensors, without necessitating solitary housing, which would cause emotional distress over the 24 weeks of this study. Besides the consideration for improving animal welfare, we also specifically do not want to model depression-linked weight effects that could be affected by isolation (Berry et al., 2012; Lander et al., 2017; Lee et al., 2020; Schipper et al., 2018). Furthermore, the difference in activity and wheel usage between single-housed and dual-housed mice is approximately known from prior literature: dual-housed mice use wheels ∼1.5x as much as single-housed mic, and dual-housed mice are ∼1.8x as active in the home cage (Zara Kanold-Tso, 2022). All animals were fed Harlan Teklad 2018 “chow” diet from weaning until 8 weeks of age, at which point they started on our HF, Harlan Teklad TD.06414. Mice in exploratory cohorts 1, 2, and 3 were all female. Mice in the validation cohort, cohort 4 (exclusively BXD43), were male and female. All mice were on a 12h day/night cycle from 7am to 7pm. Mice were checked by a trained technician or veterinarian daily, and the scientific team checked the mice every Tuesday morning, weighing the mice, then weighing and replacing the food to calculate weekly food intake (**Figure 1D**). Twelve mice died during the study, 4 of those were males of cohort 4 that were excluded for the analysis due to their heigh fighting occurrences. For one cage, both mice died shortly apart and data from that cage was suppressed, and in six other cages, one of the two died during the study. Those solitary mice were authorized to complete the study, and their data are included in all analyses. They are flagged as potential confounding factors in the Supplementary Table S2, but as only 4 “solitary” cages out of 80, their inclusion or suppression does not affect any overall conclusions.

To test whether exercise-associated gut microbiome alterations causally contribute to the metabolic phenotype observed in running versus sedentary mice, we established a fifth cohort using the same BXD43 strain, age range (8–32 weeks), running-wheel/HCM cage design, and HF diet as in Cohorts 1–4. However, in this cohort, soiled bedding was reciprocally exchanged between paired running and sedentary cages once per week to promote cross-transfer of the cage-associated microbiota. Approximately 0.5 L of soiled bedding material was transferred from each running cage to its paired sedentary cage and vice versa. All other husbandry, feeding, weighing, and IPGTT procedures followed those described above for Cohorts 1–4 (**Figure S5 A**). Following the acclimatization period, mice were switched to the HF diet at week 9, and reciprocal bedding exchange was initiated one week later, at week 10, and continued weekly thereafter. Fecal samples were collected weekly from all mice throughout the study for downstream microbiome analysis. Samples collected in weeks 8, 9, 11, 13, 16, 19, and 24 were selected for shotgun metagenomic sequencing.

In parallel, a small pilot experiment was conducted to assess the effectiveness of the bedding-exchange procedure in transferring gut microbiome alterations between cages. Sedentary BXD43 mice were maintained either on CD or HF diet, and soiled bedding was reciprocally exchanged between cages assigned to the two dietary conditions using the same procedure. Diet was selected as the microbiome-modifying factor for this validation experiment because dietary composition, particularly high-fat feeding, has been shown to exert a pronounced and reproducible effect on gut microbiome composition (Zhou et al., 2026). This pilot therefore served as a positive-control experiment to determine whether the bedding-exchange protocol was sufficient to promote cross-transfer of experimentally induced microbiome differences. All the ibformation regarding the comoposition of the different Cohort can be found in Table S2.

### Dietary Information

Our study was partially confounded by changes in the production of the Harlan Teklad 06414 high-fat diet over the course of the experiment. Although the diet was obtained from the same provider and retained the same reference number throughout the study, it was not feasible to purchase and store the entire amount required for the approximately three-year study period at once. Consequently, three separate batches of diet were ordered. All batches provided 60% of calories from fat; however, differences in their composition and physical properties were observed between batches. The first batch contained predominantly vegetable-derived lipids and was supplied as relatively firm, square pellets. Because of the more liquid nature of these lipid sources, some lipid accumulation was observed at the bottom of the food containers. In the second and third batches, a larger proportion of the lipid component was derived from lard, and the diet was supplied as softer pellets. Cohort 1 was fed exclusively with batch 1, whereas in Cohort 2 the diet batch had to be changed during the experiment. All subsequent cohorts were fed diets from batches 2 and 3. These changes in lipid composition and pellet characteristics therefore represent a potential batch effect that should be considered when interpreting differences between experimental cohorts studied at different time points.

### Home Cage Monitoring (HCM)

Home-cage activity was continuously and non-invasively monitored using the DVC system (Tecniplast S.p.A.), which detects animal movement through changes in the capacitance of an electric field generated by an array of electrodes positioned beneath each cage (Iannello, 2019). Cohorts 1–3 were housed in the earlier-generation DVC® Green Line system, whereas Cohorts 4 and 5 were housed in the newer DVC® for Emerald system. DVC cage changes were routinely performed every three weeks on Mondays by the animal caretakers. For Cohort 5, cage changes were instead performed by the experimental team on the same day as the reciprocal bedding exchange, body-weight measurements, and fecal sample collection. Home-cage locomotion data were retrieved from the Tecniplast DVC Analytics web platform. For each cage, the complete experimental period, from the first to the last day of the experiment, was selected for data extraction. The smoothed Animal Locomotion Index was selected as the locomotion activity measure, using a temporal aggregation interval of 10 min.

### Exercise Wheel Monitoring

The running mice were housed in cages in a separate sound-isolated and ventilated cabinet located in the same room as the DVC racks. The system consisted of eight individually monitored cages containing each a running wheel (Columbus Instruments; 4-inch-diameter solid plastic wheels). Wheel activity was recorded using the Columbus Instruments CI Multi-Device Interface software, with the total number of wheel revolutions continuously recorded and aggregated into 10-min intervals throughout the study. Occasional software crashes resulted in the loss of 35 days of recording in Cohort 1, 7 days in Cohort 3, 2 days in Cohort 4, and approximately 1 week in Cohort 5. Importantly, these interruptions affected only data acquisition; mice retained unrestricted access to functional running wheels throughout these periods. Running-wheel cages were smaller than the DVC cages and were not connected to an individual cage ventilation system. They also contained less bedding than the DVC cages. This reduced bedding volume was implemented following observations during pilot experiments that mice could accumulate bedding adjacent to the running wheel, potentially interfering with its movement. For Cohorts 1–4, running-wheel cages were routinely changed every two weeks. In Cohort 5, running-wheel cages were instead changed weekly by the experimental team as part of the reciprocal bedding-exchange protocol. At each cage change, a portion of soiled bedding from the corresponding paired sedentary cage was introduced into the new running-wheel cage.

### Intraperitoneal Glucose Tolerance Test

Prior to the IPGTT, mice were transferred to clean cages and fasted overnight from approximately 6 pm to 9 am. Body weight was recorded before and after fasting, and fasting blood glucose was measured using a handheld glucometer with an upper detection limit of 600 mg/dL. After fasting glucose measurements were completed, all mice received an intraperitoneal injection of 5 µL/g body weight of a 40% glucose solution prepared in 0.9% sodium chloride and pre-warmed to approximately 37°C. Blood glucose was measured at 15, 30, 45, 60, 90, and 120 min after glucose administration using blood obtained by pinprick from the distal end of the tail. Following the final measurement at 120 min, EMLA cream was applied to the tail as a topical local anesthetic, and mice were returned to their home cages with access to HF diet and water.

### Sacrifice and Tissue Collection

At the end of the study, at approximately 31 weeks of age, mice were transferred to clean cages and fasted overnight from approximately 6 pm to 9 am. Mice were then anesthetized by intraperitoneal injection of a 10:1 ketamine:xylazine mixture, corresponding to doses of 160 mg/kg ketamine and 16 mg/kg xylazine. Once fully anesthetized, mice underwent cardiac perfusion with PBS for terminal euthanasia and to remove blood from the tissues prior to collection. Perfusion was continued until the organs appeared pale, indicating adequate clearance from blood. Cecal tissue, together with several other tissues intended for subsequent studies, was then collected. Tissue collection took approximately 10 min per mouse. For Cohorts 1-4, all mice within a given cohort were sacrificed using the same procedure over a period of approximately 3-4 h. Cohort 5, which comprised 45 mice including those from the CD/HF pilot experiment, was sacrificed over two consecutive days using the same procedure. The samples have been flash frozen in dry ice upon collection and stored in the -80°C freezer.

### Microbiome Analysis

#### DNA extraction and quality control via qPCR

Frozen cecal and fecal samples were ground to a fine powder using a precooled mortar and pestle under liquid nitrogen to prevent tissue thawing. Approximately 30 mg of ground tissue was used for DNA extraction. Genomic DNA was extracted using the Wizard® Genomic DNA Purification Kit (Promega; catalog no. A1120) in combination with the Wizard® DNA Clean-Up System (Promega; catalog no. A7280), following the manufacturer’s protocol with additional modifications to improve the lysis of hard-to-lyse bacteria and facilitate the removal of food residues, as previously described (Zhou et al., 2026). Following the addition of 480 µL of cold (−4 °C) 50 mM ethylenediaminetetraacetic acid (EDTA), samples were further homogenized using five 2-mm stainless-steel beads with the Cryolys Evolution system (Bertin Technologies) for 4x90s at 6000 rpm with breaks of 30 min. The incubation following homogenization was extended to 1 h. An additional centrifugation step (2 min at 500 × g) was subsequently performed to pellet and remove residual food material. Prior to purification using the Wizard® DNA Clean-Up System, DNA concentration was determined using a NanoDrop spectrophotometer (NanoDrop 2000 C, Thermo Scientific). The volume of unpurified DNA solution loaded onto the purification column was adjusted according to the measured DNA concentration and the 20,000-ng binding capacity of the resin to optimize the purity and concentration of the recovered DNA.

The relative abundance of bacterial DNA compared with host DNA was assessed by quantitative PCR (qPCR) for all cecal samples, as previously described (Zhou et al., 2026). Two primer sets were used. Host DNA was quantified by amplification of the mouse *Gapdh* gene using GAPDH-F-2 (5′-CAAGGAGACCTCAAGGTCAG-3′) and GAPDH-R-2 (5′-GTATGCACCTCACAACGCCATG-3′). Bacterial DNA was quantified using universal prokaryotic 16S rRNA primers 926F (5′-AAACTCAAAKGAATTGACGG-3′) and 1062R (5′-CTCACRRCACGAGCTGAC-3′). Prior to qPCR, DNA concentrations were adjusted to 10 ng/µL. Reactions were performed using a LightCycler® 480 Instrument II (Roche). Each 10-µL reaction contained 2.5 µL of DNA template, 2.5 µL of a 2 µM primer mixture containing the corresponding forward and reverse primers, and 5 µL of iQ™ SYBR® Green Supermix (Bio-Rad). The amplification protocol consisted of an initial denaturation at 95 °C for 10 min, followed by 40 cycles of denaturation at 95 °C for 15 s and annealing/extension at 60 °C for 1 min. Following amplification an additional colling step at 20 °C for 1 h has been added. For each sample, ΔCp was calculated from the difference between the mean Cp values obtained with the host *Gapdh* and universal prokaryotic 16S rRNA primer sets. ΔΔCp was subsequently calculated relative to a control consisting of 60% mouse DNA and 40% microbial DNA. Relative bacterial DNA abundance was calculated as 2^ΔΔCp × 100. Samples with values >100% were considered to contain sufficient microbial DNA relative to host DNA and were selected for sequencing, whereas samples below this threshold were excluded (Zhou et al., 2026). This selection step was particularly important because samples were subjected to shotgun metagenomic sequencing rather than targeted 16S sequencing. Consequently, host DNA present in the samples would also be sequenced, reducing the proportion of sequencing reads derived from the microbial community and unnecessarily consuming sequencing capacity. As the mouse strains used in this study had already been extensively genotyped and phenotyped (Ashbrook et al., 2021), additional sequencing of host DNA was not informative for the objectives our analysis. Selecting samples with a sufficiently high microbial-to-host DNA ratio therefore maximized the sequencing depth available for microbial characterization while limiting unnecessary sequencing costs.

#### Meta genome sequencing

Metagenomic sequencing was performed by Novogene using the NovaSeq X Plus platform with 2 × 150-bp paired-end reads. Sequencing was performed with a quality specification of Q30 ≥85%, generating over 10 Gb of raw data per sample.

#### Raw data prepossessing

Raw shotgun metagenomic sequencing data were pre-processed on the high-performance computer (HPC) cluster using `KeadData v0.12.0` (Beghini et al., 2021) following the preprocessing workflow described by Zhou et al. (2026). The pipeline first performed sequence trimming and quality filtering using `Trimmomatic v0.3.9`. (Beier et al., 2026; Bolger et al., 2014), with the `KeadData` default parameter `(DEFAULT: ILLUMINACLIP:/TruSeq3-SE.fa:2:30:10 SLIDINGWINDOW:4:20 MINLEN:50)`.

Briefly, Illumina TruSeq adapter sequences are removed, reads with average base quality within a four-nucleotide sliding window below Q20 were trimmed, and post-trimming reads shorter than 50 bp were discarded. KeadData additionally uses Tandem Repeat Finder v4.09.1(Benson, 1999) to identify and discard tandem repeats. Finally, it runs `Bowtie2 v2.5.4` (Langmead & Salzberg, 2012) to remove host DNA sequences by aligning them against the C57BL/6NJ mouse reference database `(mouse_C57BL_6NJ_Bowtie2_v0.1; Huttenhower Lab)`. Following quality control and host-red depletion, the remaining sequences were used for taxonomic profiles using `MetaPlAn v4.1.1` with the `mpa_ vOct22_CHOCOPhlAnSGB_202403` database (Blanco-Miguez et al., 2023). Finally functional profiling of the metagenomic sequences was performed using the `HUMAnN 4.0.0.alpha.1` pipeline (Beghini et al., 2021), which maps reads to UniRef90 protein families (Suzek et al., 2015) and uses the resulting gene-family profiles to reconstruct and quantify metabolic pathways based on MetaCyc pathway definitions (Caspi et al., 2014).

#### Bioinformatics and Statistics phenotype

All bioinformatic and statistical analyses of taxonomic abundance data (**Table S5** and **S8**) generated with `MetaPhlAn` and pathway abundance data (**Table S6** and **S9**) generated with `HUMAnN` were performed using `R v4.5.2.` Cecal DNA samples were extracted and sequenced in three different batches (**Table S2**); therefore, potential batch effects and technical variability in the metagenomic dataset were assessed prior to downstream analyses. To determine whether technical and biological replicates exhibited the lowest dissimilarities and clustered together, Bray–Curtis dissimilarities were calculated using the `vegdist` function from the R package `vegan 2.7-6`. The resulting dissimilarity matrix was visualized as a heatmap using the R package `pheatmap 1.0.13`, with samples hierarchically clustered based on Bray–Curtis dissimilarities using average-linkage clustering. Technical variability was additionally evaluated performing analysis of variance ANOVA (`aov` function in `stats 4.5.2` package) on log10-transformed species abundances [log10(x + 1e-6)] to estimate the proportion of variance explained by biological replicates, technical replicates, their interaction, and randomly selected samples. Cohorts 3 and 4 were combined for the subsequent analyses because they followed the same experimental design and included only BXD43 mice.

For exploratory microbiome analyses, taxonomic profiles generated using MetaPhlAn 4 were analyzed at the genus and species levels. Technical replicates were excluded prior to analysis. Differences in genus-level relative abundance between Sed and Run groups were assessed using the Wilcoxon rank-sum test. Principal coordinate analysis (PCoA) was performed at the species level to assess differences in overall microbial community composition between experimental groups. Species-level relative abundances were subjected to a log1p transformation [log (x + 1)] to increase the relative contribution of lower-abundance taxa to community-level differences (McKnight et al., 2019). Bray–Curtis dissimilarities were then calculated using the `vegdist` function from the `vegan 2.7-6` R package, and PCoA was performed on the resulting dissimilarity matrix using classical multidimensional scaling `cmdscale` (Package `stats 4.5.2` in R) and Permutational Multivariate Analysis of Variance PERMANOVA has been computed using the function `adonis2` of the `vegan 2.7-6` R Package .

Differential abundance analysis was performed at the species level using the maaslin3 `1.5.1.` R package (Nickols et al., 2026), which uses multivariable models designed for sparse, compositional microbiome data. Sed and Run mice were compared within each sequencing batch, with experimental group included as a fixed effect. Prior to model fitting, taxonomic abundances were normalized within `MaAsLin3` using total-sum scaling (TSS) and subsequently log-transformed. Taxa were retained at a minimum abundance of 0.0001 and a minimum prevalence of 10%. Multiple-testing correction was performed using the built-in Benjamini–Hochberg BH procedure. `MaAsLin3` results were visualized as volcano plots using ggplot2, displaying β-coefficients against −log10(p-values); species with p < 0.05 and |β| ≥ 1 were highlighted. The same differential abundance analysis was performed for the parallel CD versus HF pilot experiment conducted alongside Cohort 5, with CD specified as the reference group.

The same procedure was applied to fecal metagenomic samples at each sampling week separately, except weeks 13 and 16 because of the reduced sample size, for Sed versus Run comparisons. Species with a joint FDR-adjusted q-value < 0.35 at at least one timepoint were visualized in bubble plots displaying MaAsLin3 abundance and prevalence effects across time. Significant associations q-value < 0.25 were observed only at week 19.

Longitudinal changes in fecal microbiome composition were additionally assessed using `MaAsLin3` with the same parameters by comparing consecutive sampling timepoints separately within the Sed and Run groups. Pairwise comparisons were performed between weeks 8–9, 9–11, 11–13, 13–16, 16–19, and 19–31. Species with an FDR-adjusted q-value < 0.25 in at least one consecutive-week comparison were retained for visualization in longitudinal heatmaps.

Similar analyses were performed on HUMAnN4 functional profiles to assess differential pathway abundance between experimental groups. Analyses were conducted both on overall pathway abundances and on species-stratified pathway profiles to determine the contribution of individual microbial species to each pathway.

All plots have been generated in R using the packages ggplot2 4.0.3, plots 3.3.0, vioplot 0.5.1, and pheatmap 1.0.13, and embellished using additional packages like RColorBrewer 1.1-3, ggalluvial 0.12.6 and scales 1.4.0..

#### Bioinformatics and Statistics phenotype

All bioinformatic and statistical analyses were performed using `R v4.5.2.`. For running and locomotion analyses, DVC locomotion-index values recorded at 10-min intervals were summed within each experimental week to obtain weekly home-cage locomotion values. For the day–night analysis, locomotion-index values were summed separately over the daytime period (07:00–19:00) and nighttime period (19:00–07:00). Running-wheel activity was similarly aggregated to obtain weekly running distances. Wheel activity, originally recorded as the number of revolutions, was converted to distance by multiplying the number of revolutions by the circumference of the running wheel, with distances subsequently expressed in kilometers. To compare longitudinal body weight, food intake, and glycemia, between experimental groups, the area under the curve (AUC) over the entire experimental period was calculated for each animal(n) respectively for each cage(n/2). AUCs were computed using the trapezoidal method via the function `trapz` within the R package `pracma 2.4.6` .

Prior to statistical testing, data normality was assessed using the Shapiro–Wilk test (`shapiro.test` in the R package `stats 4.5.2`). Comparisons between two groups were performed using Welch’s t-test (`t.test` in the R package `stats 4.5.2`) for normally distributed data and the Wilcoxon rank-sum test (`wilcox.test` in the R package `stats 4.5.2`) when the assumption of normality was not met. For comparisons involving more than two groups, ANOVA (`aov` in R package `stats 4.5.2`) was performed, followed by post-hoc pairwise comparisons (`TukeyHSD` in R package `stats 4.5.2`) to identify differences between individual groups. Where multiple comparisons were performed, p-values were adjusted for FDR using the Benjamini–Hochberg procedure. Linear associations presented in Figure 3 were assessed using Pearson’s correlation.

## Supporting information

IPGTT

Cohort Composition

Body Weight Data

Food Intake Data

Cecal Taxonomic Abundance Table

Cecal Pathway Abundance Table

Sample Information F Fecal Metagenome Samples

Fecal Taxonomic Abundance Table

Fecal Pathway Abundance Table

## SUPPLEMENTARY INFORMATION

Raw data from the DVC system and running-wheel activity as well as the complete R code are available on the GitLab repository https://gitlab.com/uniluxembourg/lcsb/gene-expression-and-metabolism/exercising_mice. Raw cecal and fecal metagenomic sequencing data are available in the NCBI BioProject repository under accession numbers PRJNA1445546 and PRJNA1242136, respectively. All other data, including body weight, IPGTT, food intake, taxonomic abundance, and pathway abundance data, are provided in Supplementary Tables S1–S9.

## ACKNOWLEDGEMENTS

Thank you to Giorgio Rosati and Tecniplast for initial piloting of the DVC equipment and data access. AL was supported by the Luxembourg National Research Fund (PRIDE21/16749720/NEXTIMMUNE2). Thanks also to the lead animal facility caretakers, Julie Virolle, Cloé Ordrenneau, and Gloria Pouit, and veterinarian Jennifer Behm and Theano Andriopoulou.

## AUTHOR CONTRIBUTIONS

Initial Project Design: DC, EGW

Phenotyping: ZZ, AL, PG, EGW

Data Handling: AL, BB

Analysis: AL, ZZ, EP, BB, EGW

DNA Extraction: AL, MG

Mice Dissection: ZZ, EGW, AL, MB, TH, PG

Paper: AL, ZZ, EGW

## REFERENCES

Allen, J. M., Berg Miller, M. E., Pence, B. D., Whitlock, K., Nehra, V., Gaskins, H. R., White, B. A., Fryer, J. D., & Woods, J. A. (2015). Voluntary and forced exercise differentially alters the gut microbiome in C57BL/6J mice. J Appl Physiol (1985), 118(8), 1059–1066. 10.1152/japplphysiol.01077.2014

Alsinani, Y., Rostamkhani, F., & Shirvani, H. (2026). Exercise and the Gut Microbiome: From Mechanisms to Clinical Applications. Nutrients, 18(10). 10.3390/nu18101565

Andreux, P. A., Williams, E. G., Koutnikova, H., Houtkooper, R. H., Champy, M. F., Henry, H., Schoonjans, K., Williams, R. W., & Auwerx, J. (2012). Systems genetics of metabolism: the use of the BXD murine reference panel for multiscalar integration of traits. Cell, 150(6), 1287–1299. 10.1016/j.cell.2012.08.012

Ashbrook, D. G., Arends, D., Prins, P., Mulligan, M. K., Roy, S., Williams, E. G., Lutz, C. M., Valenzuela, A., Bohl, C. J., Ingels, J. F., McCarty, M. S., Centeno, A. G., Hager, R., Auwerx, J., Lu, L., & Williams, R. W. (2021). A platform for experimental precision medicine: The extended BXD mouse family. Cell Syst, 12(3), 235–247 e239. 10.1016/j.cels.2020.12.002

Avila, J. J., Kim, S. K., & Massett, M. P. (2017). Differences in Exercise Capacity and Responses to Training in 24 Inbred Mouse Strains. Front Physiol, 8, 974. 10.3389/fphys.2017.00974

Beghini, F., McIver, L. J., Blanco-Miguez, A., Dubois, L., Asnicar, F., Maharjan, S., Mailyan, A., Manghi, P., Scholz, M., Thomas, A. M., Valles-Colomer, M., Weingart, G., Zhang, Y., Zolfo, M., Huttenhower, C., Franzosa, E. A., & Segata, N. (2021). Integrating taxonomic, functional, and strain-level profiling of diverse microbial communities with bioBakery 3. Elife, 10. 10.7554/eLife.65088

Beier, S., Bolger, A. M., Bolger, M. E., Schwacke, R., & Usadel, B. (2026). Trimmomatic: a decade of feature-rich, high-performance NGS read preprocessing. Bioinformatics, 42(6). 10.1093/bioinformatics/btag331

Benson, G. (1999). Tandem repeats finder: a program to analyze DNA sequences. Nucleic Acids Res, 27(2), 573–580. 10.1093/nar/27.2.573

Berry, A., Bellisario, V., Capoccia, S., Tirassa, P., Calza, A., Alleva, E., & Cirulli, F. (2012). Social deprivation stress is a triggering factor for the emergence of anxiety- and depression-like behaviours and leads to reduced brain BDNF levels in C57BL/6J mice. Psychoneuroendocrinology, 37(6), 762–772. 10.1016/j.psyneuen.2011.09.007

Blanco-Miguez, A., Beghini, F., Cumbo, F., McIver, L. J., Thompson, K. N., Zolfo, M., Manghi, P., Dubois, L., Huang, K. D., Thomas, A. M., Nickols, W. A., Piccinno, G., Piperni, E., Puncochar, M., Valles-Colomer, M., Tett, A., Giordano, F., Davies, R., Wolf, J.,…Segata, N. (2023). Extending and improving metagenomic taxonomic profiling with uncharacterized species using MetaPhlAn 4. Nat Biotechnol, 41(11), 1633–1644. 10.1038/s41587-023-01688-w

Bolger, A. M., Lohse, M., & Usadel, B. (2014). Trimmomatic: a flexible trimmer for Illumina sequence data. Bioinformatics, 30(15), 2114–2120. 10.1093/bioinformatics/btu170

Cani, P. D., Van Hul, M., Lefort, C., Depommier, C., Rastelli, M., & Everard, A. (2019). Microbial regulation of organismal energy homeostasis. Nat Metab, 1(1), 34–46. 10.1038/s42255-018-0017-4

Caspi, R., Altman, T., Billington, R., Dreher, K., Foerster, H., Fulcher, C. A., Holland, T. A., Keseler, I. M., Kothari, A., Kubo, A., Krummenacker, M., Latendresse, M., Mueller, L. A., Ong, Q., Paley, S., Subhraveti, P., Weaver, D. S., Weerasinghe, D., Zhang, P., & Karp, P. D. (2014). The MetaCyc database of metabolic pathways and enzymes and the BioCyc collection of Pathway/Genome Databases. Nucleic Acids Res, 42(Database issue), D459–471. 10.1093/nar/gkt1103

Chang, G. R., Hou, P. H., Chen, W. K., Lin, C. T., Tsai, H. P., & Mao, F. C. (2020). Exercise Affects Blood Glucose Levels and Tissue Chromium Distribution in High-Fat Diet-Fed C57BL6 Mice. Molecules, 25(7). 10.3390/molecules25071658

Choi, J., Smith, D. M., Scafidi, S., Riddle, R. C., & Wolfgang, M. J. (2024). Carnitine palmitoyltransferase 1 facilitates fatty acid oxidation in a non-cell-autonomous manner. Cell Rep, 43(12), 115006. 10.1016/j.celrep.2024.115006

Chu, N. D., Smith, M. B., Perrotta, A. R., Kassam, Z., & Alm, E. J. (2017). Profiling Living Bacteria Informs Preparation of Fecal Microbiota Transplantations. PLoS One, 12(1), e0170922. 10.1371/journal.pone.0170922

Delhanty, P. J. D., & Visser, J. A. (2025). Navigating the Strengths and Constraints of Mouse Models in Obesity Research. Endocrinology, 166(9). 10.1210/endocr/bqaf123

Denou, E., Marcinko, K., Surette, M. G., Steinberg, G. R., & Schertzer, J. D. (2016). High-intensity exercise training increases the diversity and metabolic capacity of the mouse distal gut microbiota during diet-induced obesity. Am J Physiol Endocrinol Metab, 310(11), E982–993. 10.1152/ajpendo.00537.2015

Evans, C. C., LePard, K. J., Kwak, J. W., Stancukas, M. C., Laskowski, S., Dougherty, J., Moulton, L., Glawe, A., Wang, Y., Leone, V., Antonopoulos, D. A., Smith, D., Chang, E. B., & Ciancio, M. J. (2014). Exercise prevents weight gain and alters the gut microbiota in a mouse model of high fat diet-induced obesity. PLoS One, 9(3), e92193. 10.1371/journal.pone.0092193

Figueiredo, P. A., Powers, S. K., Ferreira, R. M., Amado, F., Appell, H. J., & Duarte, J. A. (2009). Impact of lifelong sedentary behavior on mitochondrial function of mice skeletal muscle. J Gerontol A Biol Sci Med Sci, 64(9), 927–939. 10.1093/gerona/glp066

Flack, K. D., Ufholz, K., Johnson, L., Fitzgerald, J. S., & Roemmich, J. N. (2018). Energy compensation in response to aerobic exercise training in overweight adults. Am J Physiol Regul Integr Comp Physiol, 315(4), R619–R626. 10.1152/ajpregu.00071.2018

Gheorghe, C. E., Ritz, N. L., Martin, J. A., Wardill, H. R., Cryan, J. F., & Clarke, G. (2021). Investigating causality with fecal microbiota transplantation in rodents: applications, recommendations and pitfalls. Gut Microbes, 13(1), 1941711. 10.1080/19490976.2021.1941711

Hall, K. D., & Kahan, S. (2018). Maintenance of Lost Weight and Long-Term Management of Obesity. Med Clin North Am, 102(1), 183–197. 10.1016/j.mcna.2017.08.012

Hatton-Jones, K., Cox, A. J., Peart, J. N., Headrick, J. P., & du Toit, E. F. (2022). Stress-induced body weight loss and improvements in cardiometabolic risk factors do not translate to improved myocardial ischemic tolerance in western diet-fed mice. Physiol Rep, 10(2), e15170. 10.14814/phy2.15170

Herrera-Luis, E., Benke, K., Volk, H., Ladd-Acosta, C., & Wojcik, G. L. (2024). Gene-environment interactions in human health. Nat Rev Genet, 25(11), 768–784. 10.1038/s41576-024-00731-z

Hicks, J. A., Hatzidis, A., Arruda, N. L., Gelineau, R. R., De Pina, I. M., Adams, K. W., & Seggio, J. A. (2016). Voluntary wheel-running attenuates insulin and weight gain and affects anxiety-like behaviors in C57BL6/J mice exposed to a high-fat diet. Behav Brain Res, 310, 1–10. 10.1016/j.bbr.2016.04.051

Jung, A. P., & Luthin, D. R. (2010). Wheel access does not attenuate weight gain in mice fed high-fat or high-CHO diets. Med Sci Sports Exerc, 42(2), 355–360. 10.1249/MSS.0b013e3181a6d88f

Kelly, S. A., Nehrenberg, D. L., Hua, K., Garland, T., Jr., & Pomp, D. (2011). Exercise, weight loss, and changes in body composition in mice: phenotypic relationships and genetic architecture. Physiol Genomics, 43(4), 199–212. 10.1152/physiolgenomics.00217.2010

Lai, C. Q., Parnell, L. D., Smith, C. E., Guo, T., Sayols-Baixeras, S., Aslibekyan, S., Tiwari, H. K., Irvin, M. R., Bender, C., Fei, D., Hidalgo, B., Hopkins, P. N., Absher, D. M., Province, M. A., Elosua, R., Arnett, D. K., & Ordovas, J. M. (2020). Carbohydrate and fat intake associated with risk of metabolic diseases through epigenetics of CPT1A. Am J Clin Nutr, 112(5), 1200–1211. 10.1093/ajcn/nqaa233

Lander, S. S., Linder-Shacham, D., & Gaisler-Salomon, I. (2017). Differential effects of social isolation in adolescent and adult mice on behavior and cortical gene expression. Behav Brain Res, 316, 245–254. 10.1016/j.bbr.2016.09.005

Langmead, B., & Salzberg, S. L. (2012). Fast gapped-read alignment with Bowtie 2. Nat Methods, 9(4), 357–359. 10.1038/nmeth.1923

Lee, J. S., Kang, J. Y., & Son, C. G. (2020). A Comparison of Isolation Stress and Unpredictable Chronic Mild Stress for the Establishment of Mouse Models of Depressive Disorder. Front Behav Neurosci, 14, 616389. 10.3389/fnbeh.2020.616389

Leocadio, P. C. L., Oria, R. B., Crespo-Lopez, M. E., & Alvarez-Leite, J. I. (2019). Obesity: More Than an Inflammatory, an Infectious Disease? Front Immunol, 10, 3092. 10.3389/fimmu.2019.03092

Lightfoot, J. T., Turner, M. J., Daves, M., Vordermark, A., & Kleeberger, S. R. (2004). Genetic influence on daily wheel running activity level. Physiol Genomics, 19(3), 270–276. 10.1152/physiolgenomics.00125.2004

Loos, R. J. F., & Yeo, G. S. H. (2022). The genetics of obesity: from discovery to biology. Nat Rev Genet, 23(2), 120–133. 10.1038/s41576-021-00414-z

McKnite, A. M., Perez-Munoz, M. E., Lu, L., Williams, E. G., Brewer, S., Andreux, P. A., Bastiaansen, J. W., Wang, X., Kachman, S. D., Auwerx, J., Williams, R. W., Benson, A. K., Peterson, D. A., & Ciobanu, D. C. (2012). Murine gut microbiota is defined by host genetics and modulates variation of metabolic traits. PLoS One, 7(6), e39191. 10.1371/journal.pone.0039191

Merley, A. L., Hubbard, J. S., Rendahl, A. K., Duke Boynton, F. D., & Collura Impelluso, L. (2022). Behavioral and Physiologic Effects of Dirty Bedding Exposure in Female ICR Mice. J Am Assoc Lab Anim Sci, 61(1), 42–51. 10.30802/AALAS-JAALAS-21-000060

Miyoshi, J., Leone, V., Nobutani, K., Musch, M. W., Martinez-Guryn, K., Wang, Y., Miyoshi, S., Bobe, A. M., Eren, A. M., & Chang, E. B. (2018). Minimizing confounders and increasing data quality in murine models for studies of the gut microbiome. PeerJ, 6, e5166. 10.7717/peerj.5166

Niiranen, L., Stenback, V., Tulppo, M., Herzig, K. H., & Makela, K. A. (2023). Interplay between Learning and Voluntary Wheel Running in Male C57BL/6NCrl Mice. Int J Mol Sci, 24(5). 10.3390/ijms24054259

Oppert, J. M., Ciangura, C., & Bellicha, A. (2023). Physical activity and exercise for weight loss and maintenance in people living with obesity. Rev Endocr Metab Disord, 24(5), 937–949. 10.1007/s11154-023-09805-5

Perez-Munoz, M. E., McKnite, A. M., Williams, E. G., Auwerx, J., Williams, R. W., Peterson, D. A., & Ciobanu, D. C. (2019). Diet modulates cecum bacterial diversity and physiological phenotypes across the BXD mouse genetic reference population. PLoS One, 14(10), e0224100. 10.1371/journal.pone.0224100

Pernold, K., Iannello, F., Low, B. E., Rigamonti, M., Rosati, G., Scavizzi, F., Wang, J., Raspa, M., Wiles, M. V., & Ulfhake, B. (2019). Towards large scale automated cage monitoring - Diurnal rhythm and impact of interventions on in-cage activity of C57BL/6J mice recorded 24/7 with a non-disrupting capacitive-based technique. PLoS One, 14(2), e0211063. 10.1371/journal.pone.0211063

Pernold, K., Rullman, E., & Ulfhake, B. (2021). Major oscillations in spontaneous home-cage activity in C57BL/6 mice housed under constant conditions. Sci Rep, 11(1), 4961. 10.1038/s41598-021-84141-9

Podlesny, D., Durdevic, M., Paramsothy, S., Kaakoush, N. O., Hogenauer, C., Gorkiewicz, G., Walter, J., & Fricke, W. F. (2022). Identification of clinical and ecological determinants of strain engraftment after fecal microbiota transplantation using metagenomics. Cell Rep Med, 3(8), 100711. 10.1016/j.xcrm.2022.100711

Ridaura, V. K., Faith, J. J., Rey, F. E., Cheng, J., Duncan, A. E., Kau, A. L., Griffin, N. W., Lombard, V., Henrissat, B., Bain, J. R., Muehlbauer, M. J., Ilkayeva, O., Semenkovich, C. F., Funai, K., Hayashi, D. K., Lyle, B. J., Martini, M. C., Ursell, L. K., Clemente, J. C.,…Gordon, J. I. (2013). Gut microbiota from twins discordant for obesity modulate metabolism in mice. Science, 341(6150), 1241214. 10.1126/science.1241214

Roy, S., Sleiman, M. B., Jha, P., Ingels, J. F., Chapman, C. J., McCarty, M. S., Ziebarth, J. D., Hook, M., Sun, A., Zhao, W., Huang, J., Neuner, S. M., Wilmott, L. A., Shapaker, T. M., Centeno, A. G., Ashbrook, D. G., Mulligan, M. K., Kaczorowski, C. C., Makowski, L.,…Williams, R. W. (2021). Gene-by-environment modulation of lifespan and weight gain in the murine BXD family. Nat Metab, 3(9), 1217–1227. 10.1038/s42255-021-00449-w

Sanchez, P. K. M., Khazaei, M., Gatineau, E., Geravandi, S., Lupse, B., Liu, H., Dringen, R., Wojtusciszyn, A., Gilon, P., Maedler, K., & Ardestani, A. (2021). LDHA is enriched in human islet alpha cells and upregulated in type 2 diabetes. Biochem Biophys Res Commun, 568, 158–166. 10.1016/j.bbrc.2021.06.065

Sasaki, H., Hattori, Y., Ikeda, Y., Kamagata, M., Iwami, S., Yasuda, S., Tahara, Y., & Shibata, S. (2016). Forced rather than voluntary exercise entrains peripheral clocks via a corticosterone/noradrenaline increase in PER2::LUC mice. Sci Rep, 6, 27607. 10.1038/srep27607

Sasaki, T., Sugiyama, M., Kuno, M., Miyata, T., Kobayashi, T., Yasuda, Y., Onoue, T., Takagi, H., Hagiwara, D., Iwama, S., Suga, H., Banno, R., & Arima, H. (2024). Voluntary exercise suppresses inflammation and improves insulin resistance in the arcuate nucleus and ventral tegmental area in mice on a high-fat diet. Physiol Behav, 287, 114703. 10.1016/j.physbeh.2024.114703

Schipper, L., Harvey, L., van der Beek, E. M., & van Dijk, G. (2018). Home alone: a systematic review and meta-analysis on the effects of individual housing on body weight, food intake and visceral fat mass in rodents. Obes Rev, 19(5), 614–637. 10.1111/obr.12663

Stagg, D. B., Gillingham, J. R., Nelson, A. B., Lengfeld, J. E., d’Avignon, D. A., Puchalska, P., & Crawford, P. A. (2021). Diminished ketone interconversion, hepatic TCA cycle flux, and glucose production in D-beta-hydroxybutyrate dehydrogenase hepatocyte-deficient mice. Mol Metab, 53, 101269. 10.1016/j.molmet.2021.101269

Stebegg, M., Silva-Cayetano, A., Innocentin, S., Jenkins, T. P., Cantacessi, C., Gilbert, C., & Linterman, M. A. (2019). Heterochronic faecal transplantation boosts gut germinal centres in aged mice. Nat Commun, 10(1), 2443. 10.1038/s41467-019-10430-7

Strain, T., Dempsey, P. C., Wijndaele, K., Sharp, S. J., Kerrison, N., Gonzales, T. I., Li, C., Wheeler, E., Langenberg, C., Brage, S., & Wareham, N. (2023). Quantifying the Relationship Between Physical Activity Energy Expenditure and Incident Type 2 Diabetes: A Prospective Cohort Study of Device-Measured Activity in 90,096 Adults. Diabetes Care, 46(6), 1145–1155. 10.2337/dc22-1467

Suzek, B. E., Wang, Y., Huang, H., McGarvey, P. B., Wu, C. H., & UniProt, C. (2015). UniRef clusters: a comprehensive and scalable alternative for improving sequence similarity searches. Bioinformatics, 31(6), 926–932. 10.1093/bioinformatics/btu739

Sylow, L., Kleinert, M., Richter, E. A., & Jensen, T. E. (2017). Exercise-stimulated glucose uptake -regulation and implications for glycaemic control. Nat Rev Endocrinol, 13(3), 133–148. 10.1038/nrendo.2016.162

Voikar, V., & Gaburro, S. (2020). Three Pillars of Automated Home-Cage Phenotyping of Mice: Novel Findings, Refinement, and Reproducibility Based on Literature and Experience. Front Behav Neurosci, 14, 575434. 10.3389/fnbeh.2020.575434

Wang, Y., & LeCao, K. A. (2020). Managing batch effects in microbiome data. Brief Bioinform, 21(6), 1954–1970. 10.1093/bib/bbz105

Williams, A. S., Crown, S. B., Lyons, S. P., Koves, T. R., Wilson, R. J., Johnson, J. M., Slentz, D. H., Kelly, D. P., Grimsrud, P. A., Zhang, G. F., & Muoio, D. M. (2024). Ketone flux through BDH1 supports metabolic remodeling of skeletal and cardiac muscles in response to intermittent time-restricted feeding. Cell Metab, 36(2), 422–437 e428. 10.1016/j.cmet.2024.01.007

Williams, E. G., Wu, Y., Jha, P., Dubuis, S., Blattmann, P., Argmann, C. A., Houten, S. M., Amariuta, T., Wolski, W., Zamboni, N., Aebersold, R., & Auwerx, J. (2016). Systems proteomics of liver mitochondria function. Science, 352(6291), aad0189. 10.1126/science.aad0189

Yao, S., Prates, K., Freydenzon, A., Assante, G., McRae, A. F., Morris, M. J., & Youngson, N. A. (2024). Liver-specific deletion of de novo DNA methyltransferases protects against glucose intolerance in high-fat diet-fed male mice. FASEB J, 38(10), e23690. 10.1096/fj.202301546RR

Zara Kanold-Tso, D. P. (2022). Variability of wheel running behavior in mice is dependent on housing, sex, and genetic background. bioRxiv. 10.1101/2022.10.28.514233

Zhang, X., Wang, X. R., Gai, S. L., Han, Y. Q., Quan, X. Y., Yin, S., Li, J., & Wang, N. (2026). Gut microbiota reshaped by exercise improved glycolipid metabolism in obese mice via increasing the production of medium and long chain fatty acids: A multi-omics study. J Nutr Biochem, 157, 110447. 10.1016/j.jnutbio.2026.110447

Zhou, Z., Lamanna, A., Halder, R., Pansart, E., Narayanasamy, S., Boussoufa, B., Kerkour, T., Wilmes, P., & Williams, E. (2026). Integrative analysis of the mouse cecal microbiome across diet, age, and weight in the diverse BXD population. Microbiome, 14(1). 10.1186/s40168-026-02369-x

